# Alzheimer disease β-amyloid pathology accelerates age-related hearing loss: evidence from *App*^NL-F^ “knock-in” mice

**DOI:** 10.64898/2026.05.25.727095

**Authors:** Verónica Fuentes-Santamaría, Juan C. Alvarado, Zaskya Benitez, María C. Gabaldón-Ull, Takaomi C. Saido, Takashi Saito, Thomas Lenarz, José M. Juiz

## Abstract

Age-related hearing loss (ARHL) is a main acquired risk factor for dementia, including Alzheimer disease (AD), but links are unknown. We are using a mouse model with traits of both aging pathologies to test mechanistic interactions. The “knock-in” *App*^NL-F^ mouse reproduces β-amyloid pathology in brain regions homologous to those involved in human AD. Because it was generated from the C57BL/6J mouse, it expresses early signs of ARHL, previously reported in this inbred strain. We found evidence that the early-onset ARHL of the C57BL/6J mouse is accelerated in the *App*^NL-F^ mouse. In adult C57BL/6J mice around seven-month-old, there were significant increases in auditory thresholds. In adult age-matched *App*^NL-F^ mice, auditory thresholds were significantly more elevated, suggesting acceleration of ARHL. In old mice, past thirteen months of age, hearing thresholds were equally elevated in both strains. Outer hair cell loss was significantly increased in adult *App*^NL-F^ relative to age-matched C57BL/6J mice, progressing from basal to apical cochlear turns. Spiral ganglion neuron loss also was larger. In adult *App*^NL-F^ mice there was more atrophy and enlarged capillary lumen size in the stria vascularis (SV), supporting accelerated ARHL. These findings suggest that central β-amyloid pathology worsens age-related damage to the auditory receptor, thus accelerating ARHL. Damage to the SV and its capillaries in *App*^NL-F^ mice point to exacerbation of strial and vascular pathology in the aging cochlea by central β-amyloid pathology. ARHL acceleration by central β-amyloid pathology may contribute to a vicious circle with implications for prevention and therapies.

**Highlights:**

- Age-related hearing loss worsens in a mouse model of Alzheimer β-amyloid pathology.
- Hearing thresholds further increase relative to naturally occurring hearing loss.
- Loss of outer hair cells and spiral ganglion neurons is larger.
- The stria vascularis and its microcirculation are more atrophic and damaged.
- Alzheimer disease may potentiate peripheral presbycusis.

## 1. Introduction

Human age-related hearing loss (ARHL), or presbycusis, is the consequence of cumulative age-dependent damage and neurodegeneration of the auditory receptor organ and central pathways [1–3]. It has been historically viewed more as an inevitable collateral nuisance of the aging process, rather than a condition worth of systematic management in the context of systemic organic decline [4]. Although such misconception still lingers, perspective is changing [5,6]. The increase in life expectancy, particularly dramatic in the last 60 years [7], has not been sufficiently balanced by improvements in healthy aging; hence, the current impact of age-related pathologies is greater, affecting a larger fraction of the population [8]. This is the case for ARHL, one of the most frequent chronic conditions during aging and increasingly recognized as major source of aging disability [9,10]. Such growing impact of ARHL in healthy aging is fostering redefinition of priorities in its understanding and management [11].

ARHL manifests by loss of hearing sensitivity, typically progressing from higher to lower frequency components of sound, and difficult separation of auditory signals from noise with poor speech discrimination [2]. By impairing sound and speech perception, degraded auditory input in ARHL limits communication and thus potentiates social withdrawal in the elderly, which is a risk factor for morbidity and all-cause mortality, contributing to reduced cognitive and physical activity [12,13]. Thus, auditory deprivation in the elderly participates in the chain of events leading to limited cognitive performance [13]. Although this underscores by itself the growing relevance of ARHL in aging, ARHL may have more direct links with profound deterioration of the cognitive domain in the elderly. Strong associations of ARHL with dementia, particularly Alzheimer disease (AD), have been reported. A pioneering AD case-control study [14] showed that hearing loss ≥30 dB HL was more prevalent in the AD case group, and hearing loss correlated independently with the severity of cognitive dysfunction. More recent evidence added strong support to the notion that aged people with ARHL are more likely to develop AD over time than those with better hearing preservation [4,15–17]. Also, individuals with objective or self-reported hearing impairment showed an accelerated rate of brain atrophy compared to those with normal hearing [18,19]. Recent meta-analysis identified ARHL as the single major modifiable risk factor of AD [20]. It has been estimated that elimination of hearing loss represents a potential risk reduction of 7% to 9% in AD prevalence, depending on the studies included in the meta-analysis [20,21].

One hypothesis stemming from this emerging notion is that there may be mechanistic links between ARHL and AD [22]. Identifying evidence of such links may help establish biological markers for the risk of cognitive decline associated with presbycusis, thereby contributing to the optimization of preventive and therapeutic strategies for ARHL focused on cognitive impairment and AD. [23].

To unravel possible pathophysiological interactions, animal models simultaneously reproducing controlled traits of AD and ARHL are required. As far as AD is concerned, genetically modified mouse models are widely used to reproduce altered protein homeostasis and accumulation in the brain of defective protein degradation products characteristic of this disease [24]. For instance, there are numerous mice strains genetically tailored to reproduce faulty processing of the amyloid precursor protein (APP) leading to β-amyloid (Aβ) pathology, an early hallmark in the progression of AD [24–27]. “Standard” AD transgenic mice technology takes advantage of random chromosomal insertion of *App* transgenes incorporating mutations [28]. As a result, APP is overexpressed and incorporated mutations override normal APP proteolytic degradation. This leads to extracellular accumulation of Aβ peptides, with insoluble forms packed in senile plaques. These transgenic mice lines, in which mutated sequences of APP proteolytic enzymes also are incorporated, are essential tools to understand pathophysiological roles of Aβ peptides [26]. However, widespread distribution of Aβ deposits throughout the encephalon may be a confounding factor to unravel interactions between AD traits and other conditions such as ARHL, because Aβ accumulation in regions not affected in AD may hamper interpretation of results [26,29]. Gene-targeted “knock-in” (KI) mice technology [28] overcomes this limitation. One of these KI AD mouse models is the *App*^NL-F^ strain [30,31]. In this strain, the endogenous *App* gene is modified by inserting human amino acid coding sequences of the Aβ region into the mouse sequence, which is then re-inserted along with its promoter at the same locus [25,31]. Homologous recombination leads to expression of such “humanized” APP only in neurons that express this protein under physiological conditions, which are the ones primarily affected in AD. Thus, it is possible to maintain normal levels and localization of APP while reproducing the effects of modifications in its expression/regulation [24,25]. For this, the Aβ region of the humanized *App* gene construct is fitted with mutated sequences which induce proteolytic cleavage at different sites in the protein products and thus increased levels and different proportions of Aβ fragments, prone to accumulate in plaques. The proportion of Aβ types and its deposition may be manipulated for a controlled reproduction of AD traits [30,31]. Specifically, the *App*^NL-F^ mouse contains the familiar Iberian (F) and Swedish (NL) mutations in the Aβ sequence. Homozygote *App*^NL-F^ mice develop amyloid plaques at six months of age, along with inflammation and gliosis. Synaptic loss is present at nine months [30,31]. Memory impairment is evident at eighteen months [30,31]. Altogether, the *App*^NL-F^ mouse is currently regarded as a well-fitted model for sporadic AD in pre-clinical/early cognitive impairment stages when interventions are likely to be more successful [25,30].

Crucially for our aims, as far as auditory function is concerned, the *App*^NL-F^ mouse, like most other genetically modified mice strains, was generated on the genetic background of the C57BL/6J mouse [25,30,31]. This inbred mouse strain develops early hearing loss around six months of age, functionally reminiscent of human ARHL [32–35]. This includes increased auditory thresholds starting in the high frequency range and progressive loss of outer hair cells and spiral ganglion neurons [34,35]. Because of the characteristics of the hearing deficit, the C57BL/6J mouse is a convenient and generally accepted model of early onset hearing loss [36] with traits of peripheral human ARHL [32–35]. We propose that this ARHL phenotype in combination with the “AD comorbidity” provided by the “humanized” *App* gene in the *App*^NL-F^ mouse, may be a useful, so far unexploited model to search for mechanistic interactions between AD and ARHL. Current hypotheses of such possible mechanistic interactions focus on the “ascending” effects of age-related auditory deprivation on the cognitive domain and cognitive regions of the brain [22,37]. However, the possibility that the converse may also take place, i.e., that central AD pathology, specifically Aβ pathology, may affect the course of ARHL has received less attention [38,39]. The *App*^NL-F^ mice is particularly well suited to address this relevant question. Thus, using a combination of auditory brainstem response recordings and quantitative histology, we have tested in *App*^NL-F^ mice whether and how AD Aβ pathology may affect the progression of ARHL.

## 2. Materials and Methods

### 2.1. Experimental animals and design

*App*^NL-F^ and wild type (WT) C57BL/6J mice (onwards, WT C57) were used. Sample size was estimated using Boston University sample size calculator (https://www.bu.edu/research/forms-policies/iacuc-sample-size-calculations/). Statistical power was set al 95% with an β level of 0.01 for an effect size of 5 dB SPL, taking a normal hearing threshold of 35 dB SPL at 8 kHz as baseline. Calculations rendered a sample size range of 24 to 30 mice for each condition, experimental and control. Homozygous *App*^NL-F^ mice and control WT C57 mice were distributed in three age groups: “Young”, from 2 to 4 months old, (average 2.8 ± 0.07 months for WT C57 and 2.87 ± 0.09 months for *App*^NL-F^), “Adult”, from 6 to 8 months, (average 7.32 ± 0.05 and 7.30 ± 0.03 months, respectively) and “Old”, from 12 to 14 months (average 13.38 ± 0.01 and 14 ± 0.01 months respectively). Analysis was conducted in 10 animals per age group and strain, resulting in a total of sixty mice, thirty per condition, experimental and control, fitting sample size power calculations. Results were pooled across sex, because there were no observable differences between males and females (close to 50/50 in the sample).

Mice were bred at the Animal Experimentation Service of the University of Castilla-La Mancha (UCLM) in Albacete. The UCLM *App*^NL-F^ colony was established from the original one at the Riken Institute (Wako, Japan), through a Material Transfer Agreement. Breeding pairs were kindly donated by Prof. Carlos Dotti, from the Center for Molecular Biology “Severo Ochoa” at the Universidad Autónoma de Madrid (UAM). The use, care and handling of the animals followed current regulations. Procedures and protocols approved by the competent authority were supervised and monitored by UCLM’s Animal Experimentation Service.

### 2.2. Brainstem auditory evoked potential

#### 2.2.1 Auditory brainstem response recordings

Procedures were carried out in a soundproof room. Mice were anesthetized by inhalation of 4% isoflurane (flow rate of 1 L/min OR_2_) for induction, followed by injection of ketamine/xylazine (ketamine 100mg/kg and xylazine 10mg/kg) for maintenance, and placed for recording inside a cabin with acoustic and electrical insulation (Incotron Eymasa S.L., Barcelona, Spain). During recording, the body temperature was monitored and maintained at 37.5°C with a non-electric heating pad. Disposable needle electrodes (Technomed Medical Accessories, Netherlands, TE/S05723-001) were inserted subdermally on the cranial vertex (non-inverted electrode) and on the right (inverted electrode) and left (ground) mastoid processes. Auditory stimulation and recordings were performed using equipment and software from Intelligent Hearing Systems (Miami, FL, USA). Stimuli were generated digitally with SmartEP software (Intelligent Hearing Systems), using the Universal Smart Box and utility. For signal amplification, the Opti-Amp transmitter amplifier was used. High frequency responses were recorded with the High Frequency Transducer equipment, connected to the Sound Output Booster Box. Stimuli consisted of pure tones (5 ms rise/fall time without plateau, 20 times per second) at 4 different frequencies (4, 6, 8 and 16 kHz) and clicks. Stimuli were presented in the right outer ear canal using an EDC1 electrostatic speaker driver (Tucker-Davis Technologies -TDT-, Miami, FL, USA) via an EC-1 electrostatic speaker (TDT). Stimuli were calibrated prior to the experiments with SmartEP software (Intelligent Hearing Systems) and an ER-10B+ low-noise microphone (Etymotic Research Inc., Elk, Groove, IL, USA). Responses were low pass filtered (0.3-5 kHz), after which they were averaged (starting at 1024 waves) and stored for further analysis.

#### 2.2.2 Auditory thresholds and threshold shifts

To assess auditory thresholds, background activity (background noise) was measured prior to stimulus onset, and evoked responses were recorded starting at 80 dB SPL, in descending 5 dB SPL steps. Auditory threshold was considered the lower stimulus intensity that evoked waves with a voltage amplitude greater than 2 standard deviations (SD) relative to background activity (base noise).

The threshold shift was calculated for each of the frequencies studied by subtracting auditory thresholds in the adult and old age groups from the auditory thresholds in the young mice age groups.

### 2.3. Histological assessment

#### 2.3.1. Processing of the cochleae for histology

Procedures followed protocols established in the lab for rodent cochlear histology [40–42]. Briefly, after deep anesthesia with sodium pentobarbital (200mg/Kg) injected intraperitoneally, mice were perfused transcardially with 0,9% saline wash followed by a phosphate-buffered 4% paraformaldehyde (PF) for tissue fixation. After opening the skull, cochleae were detached from the surrounding bone and postfixed by intra-scalar instillation of PF, followed by immersion in the same fixative for 48 hours. After extensive washing in 0.1 M phosphate buffer (PB 0.1M; pH 7.3), cochleae were decalcified in 50% RDO rapid decalcification solution for 1h.

The right cochleae were used for cochlear whole-mount preparations which were immunostained with prestin for outer hair cell (OHC) counts, as described below. Left cochleae were cryoprotected in 30% sucrose in PB 0,1M, embedded in 15% sucrose and 10% gelatin solution and frozen in a 2-propanol/dry ice cooling bath at −70°C. Cochlear mid-modiolar sections (12µm), obtained with a cryostat, were serially mounted onto coated slides. Alternate slide sets were routinely stained with 1) Hematoxylin/eosin (H/E) for microscopy analysis, including whole stria vascularis (SV) morphometry and spiral ganglion (SG) neuron relative packing density counts (Fuentes-Santamaría et al., 2024) or 2) Biotinylated phalloidin (Pha) (1:100; B7474; Biotin-XX; ThermoFisher Scientific) which was then visualized with streptavidin conjugated to Alexa 594 (Molecular Probes, Eugene, OR, USA) for area measurements of blood vessels in the stria vascularis (SV).

#### 2.3.2. Stria vascularis and spiral ganglion neuron morphometry on cochlear sections

For comparative measurements of the thickness of the SV, H/E stained mid-modiolar cochlear sections 24 µm apart; four sections per animal, 4 mice per group, were used as reported elsewhere [41,42]. Images from selected fields were captured with a 40x objective, and SV thickness measured using Scion Image (version beta 4.0.2; developed by Scion Corp). For thickness measurements, the SV in each cochlear turn the sectional profile was divided into 10 orthogonal measurement planes along its length. Obtained length values were averaged and expressed as thickness in βm ± SEM for each cochlear turn.

Comparative survival ratios and area reduction of spiral ganglion (SG) neurons were carried out on H/E mid-modiolar cochlear sections (24 µm apart; four sections per animal, 4 animals per group). Only neurons with a well-delineated cytoplasm and visible nucleus and nucleolus were included in the analysis, as described previously [42]. Images of selected fields obtained with a 40x objective were captured, and the number of neurons/field and their cross-sectional areas measured with Scion Image. Data were normalized relative to the control condition (young animal groups) and expressed for each cochlear turn as % SG neuron survival ±SEM and % SG neuron area reduction.

#### 2.3.3. Outer hair cell counts on whole mount preparations of cochlear turns

*Prestin immunolabeling.-* As previously described, the right cochlea of each animal was removed and decalcified. The spiral turns of the organ of Corti were carefully exposed and dissected out in segments corresponding to basal, middle and apical portions for OHC counts (Alvarado et al 2020). Each cochlear turn was incubated “in toto” with anti-prestin antibody (1:100; SC-22692, Santa Cruz, Biotechnology, Inc. Germany) diluted in phosphate-buffered saline (PBS) containing 0.2% Triton X-100 (Tx) for 24h at 4°C. The next day, cochlear turns were incubated in donkey anti-goat conjugated to Alexa 594 secondary antibody for 2h, rinsed, counterstained with DAPI nuclear staining and cover slipped. Fluorescence was visualized using a confocal laser scanning microscope (LSM 710; Zeiss, Jena, Germany) equipped with a 40X Plan Apo oil-immersion objective (1.4 NA) and excitation laser lines at 405 and 594 nm. Series of z-stack confocal images (3–5 μm thickness, 1024 × 1024 pixels) were acquired at intervals of 0.5 μm and saved as TIFF files.

##### Outer hair cell counts

Cochlear surface preparations were used for OHC counts, as described previously [41]. Briefly, cell counts were performed in segments of approximately 250 μm-long along the length of the organ of Corti using the public domain image analysis software Scion Image (version beta 4.0.2; developed by Scion Corp). The specificity of prestin immunolabeling allowed unambiguous identification of present and missing OHC profiles. OHC survival was thus calculated an expressed as the percentage of remaining OHC along the length of the basilar membrane in *App*^NL-F^ mice relative to control C57.

### 2.4. Statistical tests

Data are expressed as mean ± SEM. Measurements of ABR parameters were performed at 80 dB SPL unless otherwise stated. Two-way repeated measures analysis of variance (ANOVA) was used for hypothesis testing. Statistically significant effects over age and mice strain were evaluated. If the main analysis indicated a significant effect of one factor or an interaction between factors, a Scheffé post hoc test was performed. Significance levels (α) and power (β) were set to 0.05 and 95%, respectively. As far as auditory thresholds are concerned, Pearsońs correlation analysis was carried out to test the strength of linear association between age and click auditory thresholds, as well as pure tone auditory thresholds at 4, 6, 8 and 16 kHz for each mice strain.

## 3. Results

### 3.1. Age-related changes in ABR waveforms in WT C57 and *App*^NL-F^ mice

Figures 1 and 2 show representative ABR recordings at 8 kHz from WT C57 and *App*^NL-F^ mice across the three age groups. Young mice (2–4 months) of both strains displayed the typical five-peak wave pattern characteristic of mice and other mammals [43], identifiable from 80 dB SPL down to 30 dB (Figs. 1 and 2, arrows, first column).

**Figure 1.**
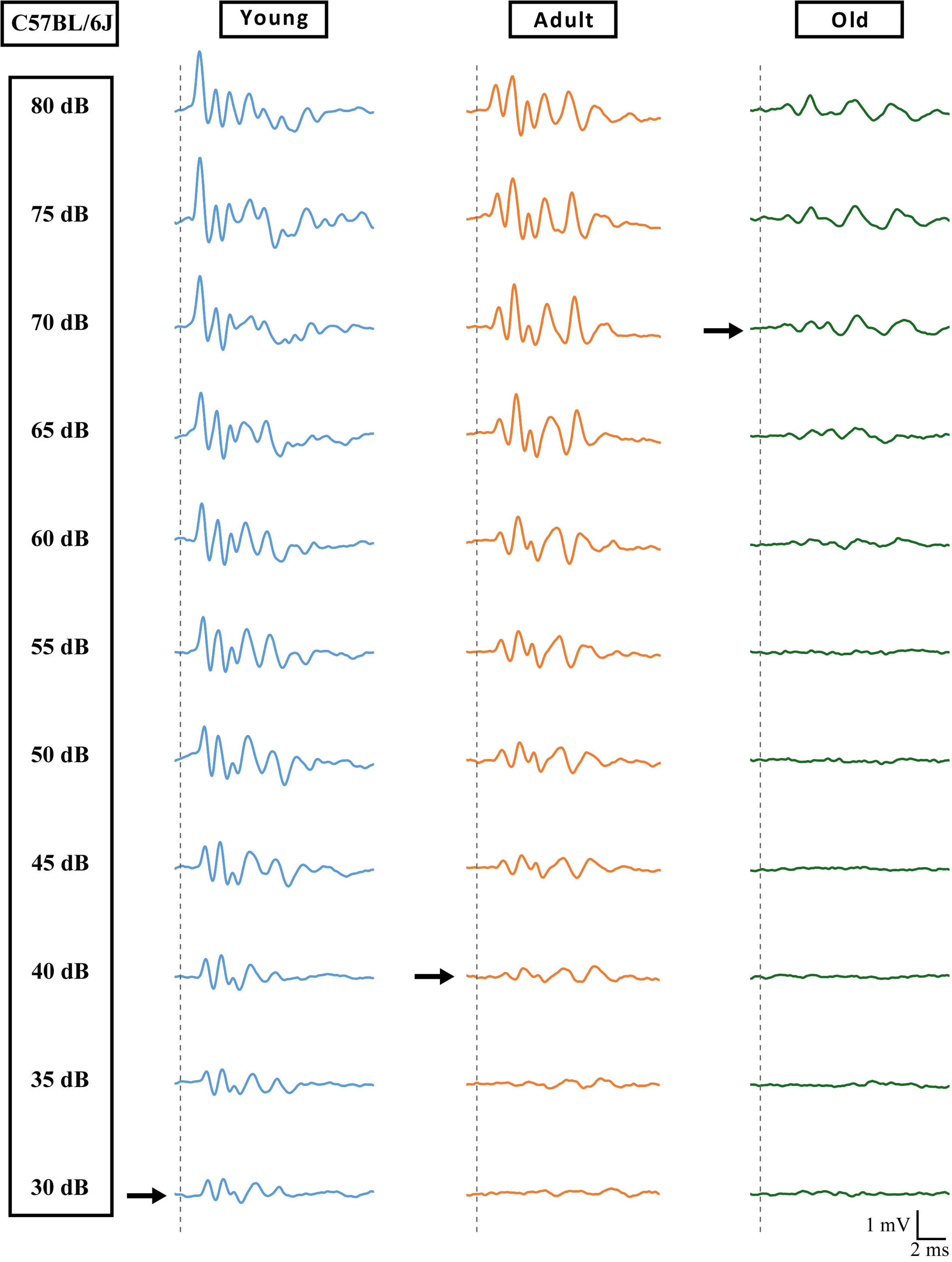
Representative ABR recordings from young, adult and old wild type C57BL/6J (WT C57 in the text), illustrating progression of ARHL. A normal wave pattern with five positive peaks is seen down to 30 dB SPL in the young mouse (black arrow, first column). In the adult, no evoked activity is detectable below 40 dB SPL (black arrow, middle column), and below 70 dB SPL in the old mouse (black arrow, third column). Dashed line indicates the stimulus onset, and the arrows show the auditory threshold.

**Figure 2.**
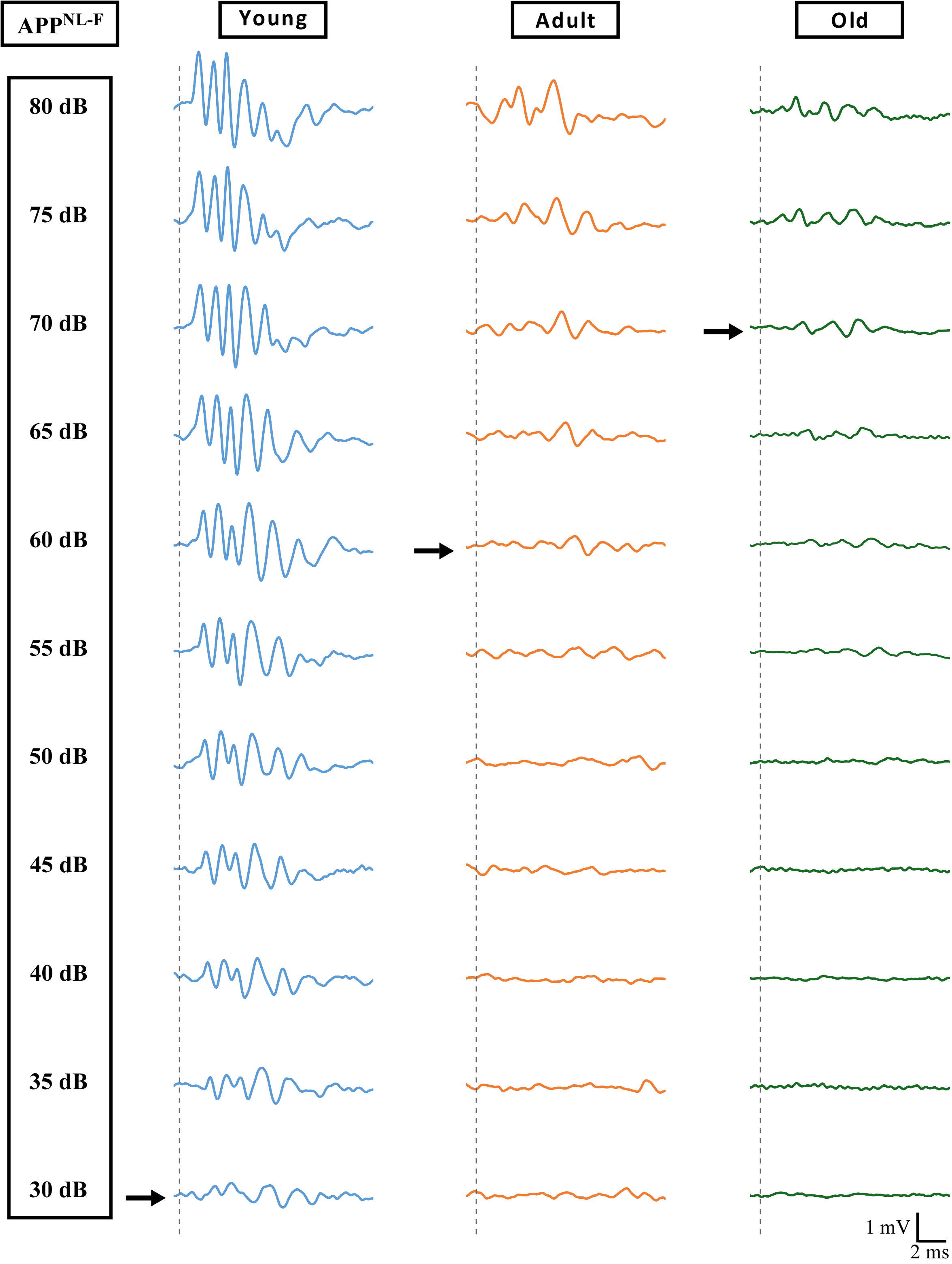
Representative ABR recordings from young, adult and old *App*^NL-F^ mice, illustrating acceleration of ARHL. Like in C57BL/6J, in the young *App*^NL-F^ mouse, a five-peak wave pattern is identifiable down to 30 dB SPL (black arrow, first column). In the adult *App*^NL-F^ mouse, no activity is detectable below 60 dB SPL (black arrow, middle column). Like in C57BL/6J, in old *App*^NL-F^ mice, no activity is recordable below 70 dB (black arrow, third column). Thus, ARHL in *App*^NL-F^ mice. Compare with Figure 1. Dashed line indicates the stimulus onset, and arrows show the auditory threshold.

In adult mice, WT C57 retained a well-defined five-wave pattern from 80 to 40 dB SPL (Fig. 1, middle column), whereas in adult *App*^NL-F^ mice waves progressively attenuated between 75 and 60 dB and were no longer recordable below 60 dB (Fig. 2, middle column). In old mice of both strains, identifiable waves were detectable only down to 70 dB SPL and amplitudes were visibly reduced relative to young and adult animals (Figs. 1 and 2, third row). Overall, ABR waveforms reflected an age-related decline in auditory sensitivity that emerged earlier in t *App*^NL-F^ than in WT C57 mice.

### 3.2. Absolute thresholds and threshold shifts reveal accelerated auditory decline in *App*^NL-F^ mice

Click and pure-tone thresholds for each strain, age group and frequency are summarized in Table 1 and Fig. 3A,B. In young WT C57 mice, click threshold averaged 36.67 ± 2.79 dB, and pure-tone thresholds followed the expected pattern of higher values at 16 kHz and lower values at 8, 6 and 4 kHz. Young *App*^NL-F^ mice did not differ significantly from age-matched young WT C57 mice, either for clicks (36.43 ± 4.04 dB) or for any pure-tone frequency (Fig. 3A,B; Table 1).

**Figure 3.**
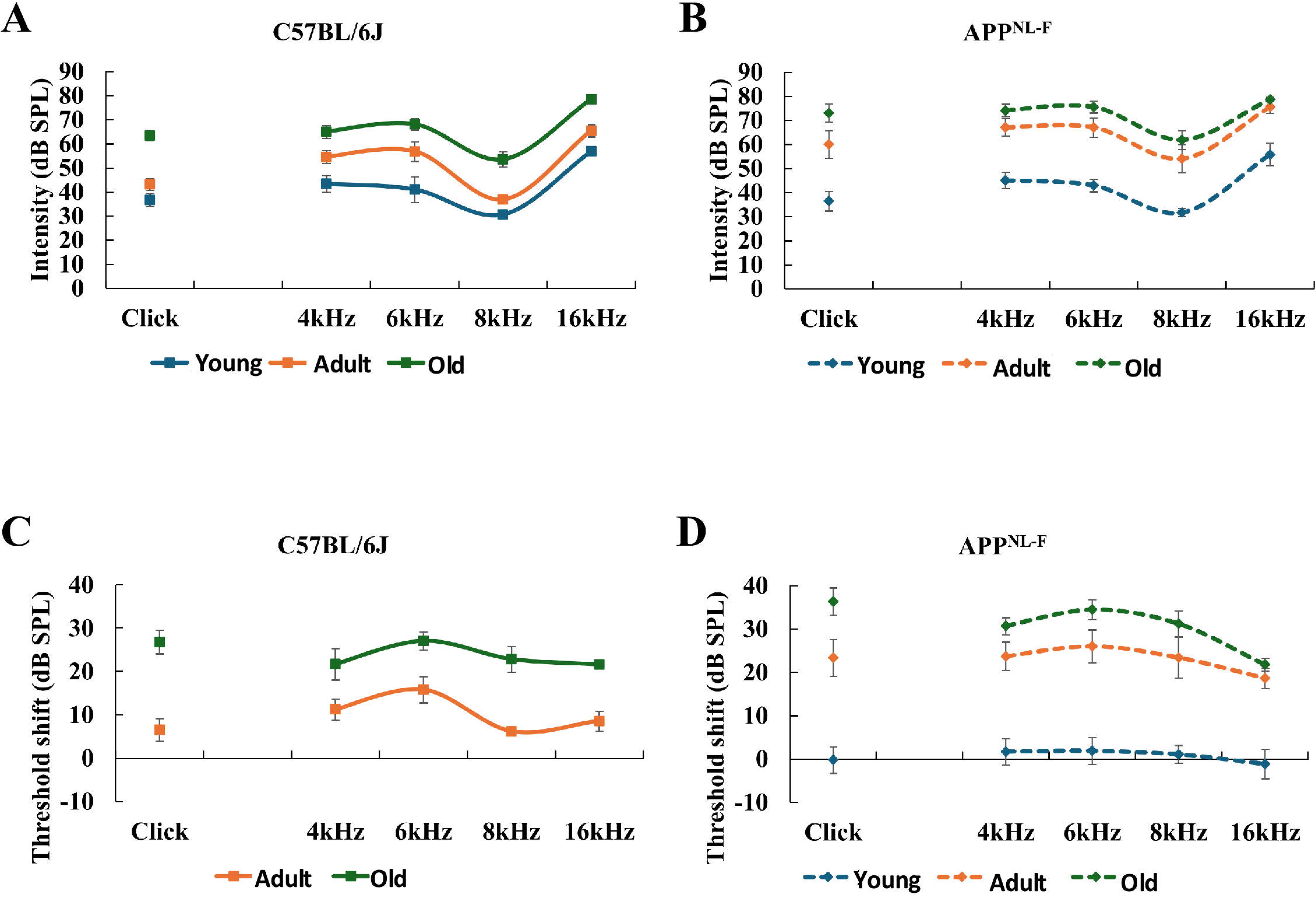
Absolute thresholds (A,B) and threshold shifts (C,D) in wild type C57BL/6J and *App*^NL-F^ mice. In C57BL/6J mice (A), click thresholds do not differ statistically between young and adults, whereas threshold elevation is significant in old mice. In *App*^NL-F^ mice (B) click thresholds increase significantly between young and adult ages. Click thresholds in adult *App* ^NL-F^ mice (B, orange square, 60 dB SPL), are close those in *old* wild type C57BL/6J mice (A, green square, 63.4 dB SPL). In young mice of both strains, frequency thresholds are similar throughout the tested frequency range (A, continuous blue curve and B, dashed blue curve). Frequency thresholds increase in adult mice of both strains, although overall the threshold curve is more elevated in adult *App*^NL-F^ mice (continuous and dashed orange lines in A and B). Frequency threshold curves are further elevated in C57BL/6J mice (orange and green curves in A). Old *App*^NL-F^ mice frequency threshold curves are comparable to old C57BL/6J, but not significantly more elevated than in adult *App*^NL-F^ mice (B, dashed orange and green curves). Thus, auditory thresholds increase faster during aging in *App*^NL-F^ mice. Threshold shifts (C, D) are larger between young and adult *App*^NL-F^ mice (D, blue and orange squares, dashed curves) than between their C57BL/6J counterparts (C, baseline and orange squares, continuous curve). Curves are closer between adult and old *App*^NL-F^ mice than between adult and old wild type C57BL/6J (C, D, orange and green), further supporting accelerated ARHL in *App*^NL-F^ mice. See Table 1 for statistics details.

**Table 1.**
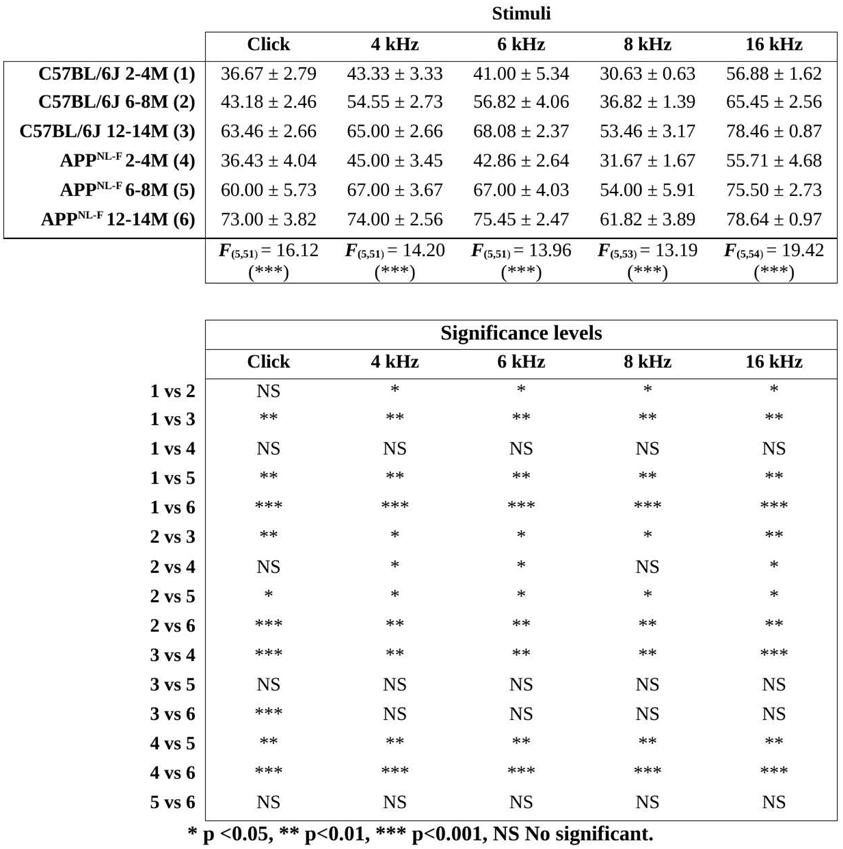
Mean ± SEM and ANOVA of the interaction between age and strain over auditory thresholds.

In adult WT C57 mice, click threshold rose to 43.18 ± 2.46 dB, a non-significant increase over the young group, whereas pure-tone thresholds were significantly elevated at all frequencies (Fig. 3A; Table 1). In adult *App*^NL-F^ mice, click threshold averaged 60.00 ± 5.73 dB, a statistically significant increase relative to age-matched adult WT C57 and to young mice. Pure-tone thresholds were likewise significantly raised across frequencies, producing an elevated and flattened threshold curve (Fig. 3A,B; Table 1).

In old WT C57 mice, click threshold further rose to 63.46 ± 2.66 dB, a significant increase over the adult group, and pure-tone thresholds at all frequencies were also significantly elevated (Fig. 3A; Table 1). In old *App*^NL-F^ mice, click threshold (73.00 ± 3.82 dB) was significantly higher than in old WT C57 and in adult WT C57 mice, although the additional rise over adult *App*^NL-F^ did not reach statistical significance (Fig. 3B; Table 1). Pure-tone thresholds in old *App*^NL-F^ mice exceeded those of old WT C57 at frequencies below 16 kHz, but neither these differences nor those with adult *App*^NL-F^ mice were significant (Fig. 3B; Table 1).

Threshold shifts relative to young WT C57 mice (Fig. 3C,D) confirmed this pattern. Adult WT C57 mice showed modest shifts (6.52 ± 2.62 dB for clicks; 6.19–15.82 dB across pure tones), which increased markedly in old WT C57 mice (26.79 ± 2.72 dB for clicks; 21.59–27.08 dB for pure tones). In *App*^NL-F^ mice, shifts were essentially absent in the young group (−0.24 ± 3.06 dB for clicks; −1.16 to 1.86 dB for pure tones) but large in the adult group (23.33 ± 4.24 dB for clicks; 18.63–26.00 dB for pure tones) — that is, of comparable magnitude to the shifts seen only at old age in WT C57 mice. Old *App*^NL-F^ mice displayed the largest shifts (36.33 ± 1.96 dB for clicks; 21.76–34.45 dB for pure tones). Taken together, threshold-shift analysis indicates an earlier onset and larger magnitude of auditory decline in *App*^NL-F^ than in WT C57 mice.

Correlation analysis between age and thresholds (Fig. 4) showed strong positive associations in both strains. Pearson’s r was higher in WT C57 than in *App*^NL-F^ mice for clicks (0.85 vs. 0.68) and for the two highest frequencies (8 kHz: 0.84 vs. 0.58; 16 kHz: 0.85 vs. 0.63), and similar between strains at 4 kHz (0.72 vs. 0.70) and 6 kHz (0.70 vs. 0.73). The flatter regression slopes and lower r values for clicks and high frequencies in *App* ^NL-F^ mice reflect their early threshold rise followed by a plateau at old age, consistent with accelerated ARHL.

**Figure 4.**
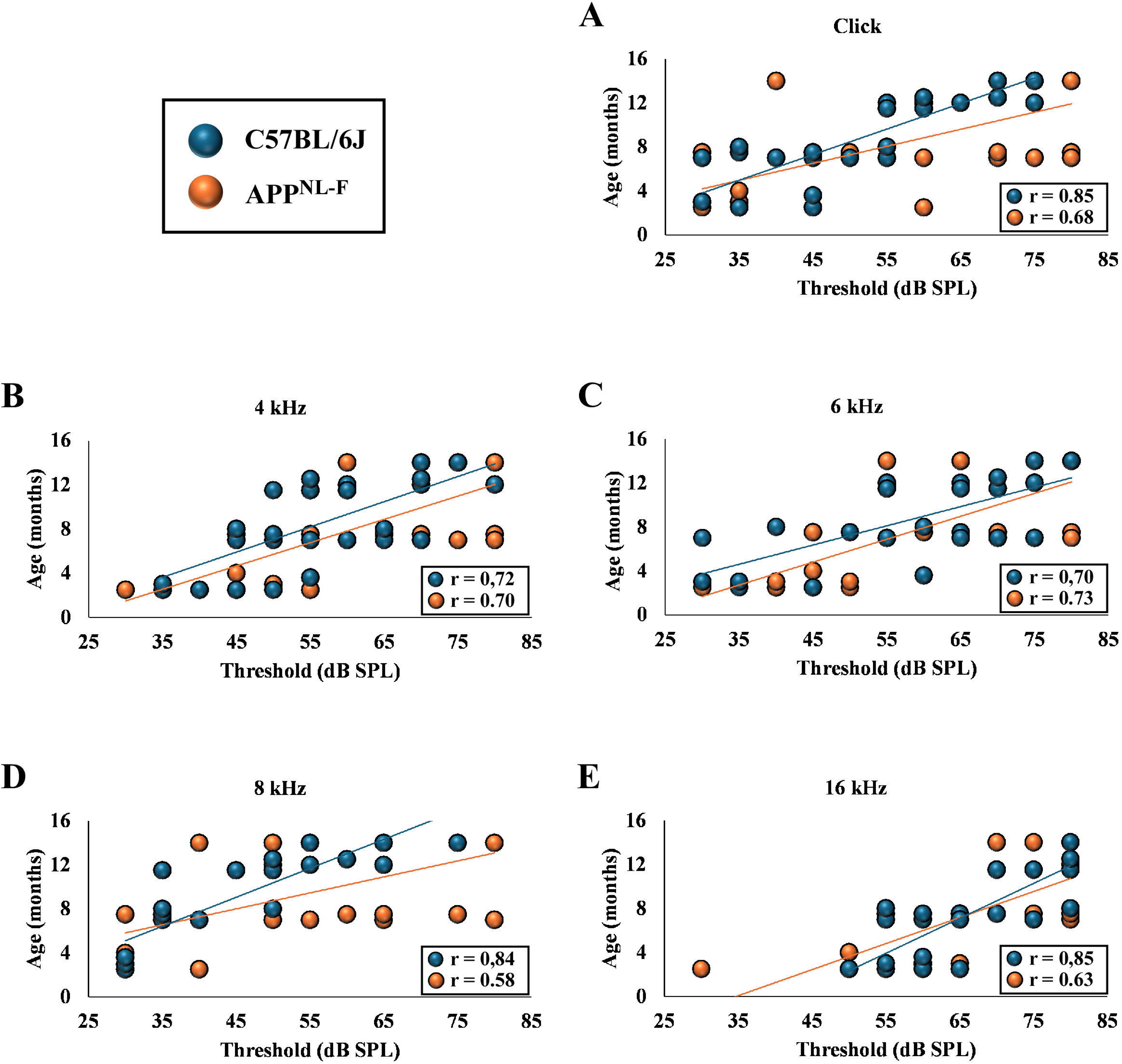
Regression plots and correlation between age and auditory thresholds (A: clicks, B, C, D, E: tonal frequencies) in wild type C57BL/6J and *App*^NL-F^ mice. Smaller Pearsońs correlation coefficient (r) values and flatter regression lines in *App*^NL-F^ mice for clicks (A) and for the highest tested frequencies of 8 kHz (D) and 16 kHz (E) point to accelerated threshold increases.

### 3.3. Accelerated outer hair cell loss in *App*^NL-F^ mice

Whole-mount preparations of cochlear turns labelled *in toto* with prestin immunofluorescence showed staining restricted to the three rows of outer hair cells (OHCs) (Fig. 5); prestin immunolabelling of the OHC plasma membrane allowed unambiguous identification of individual sensory cells (e.g., Fig. 5H).

**Figure 5.**
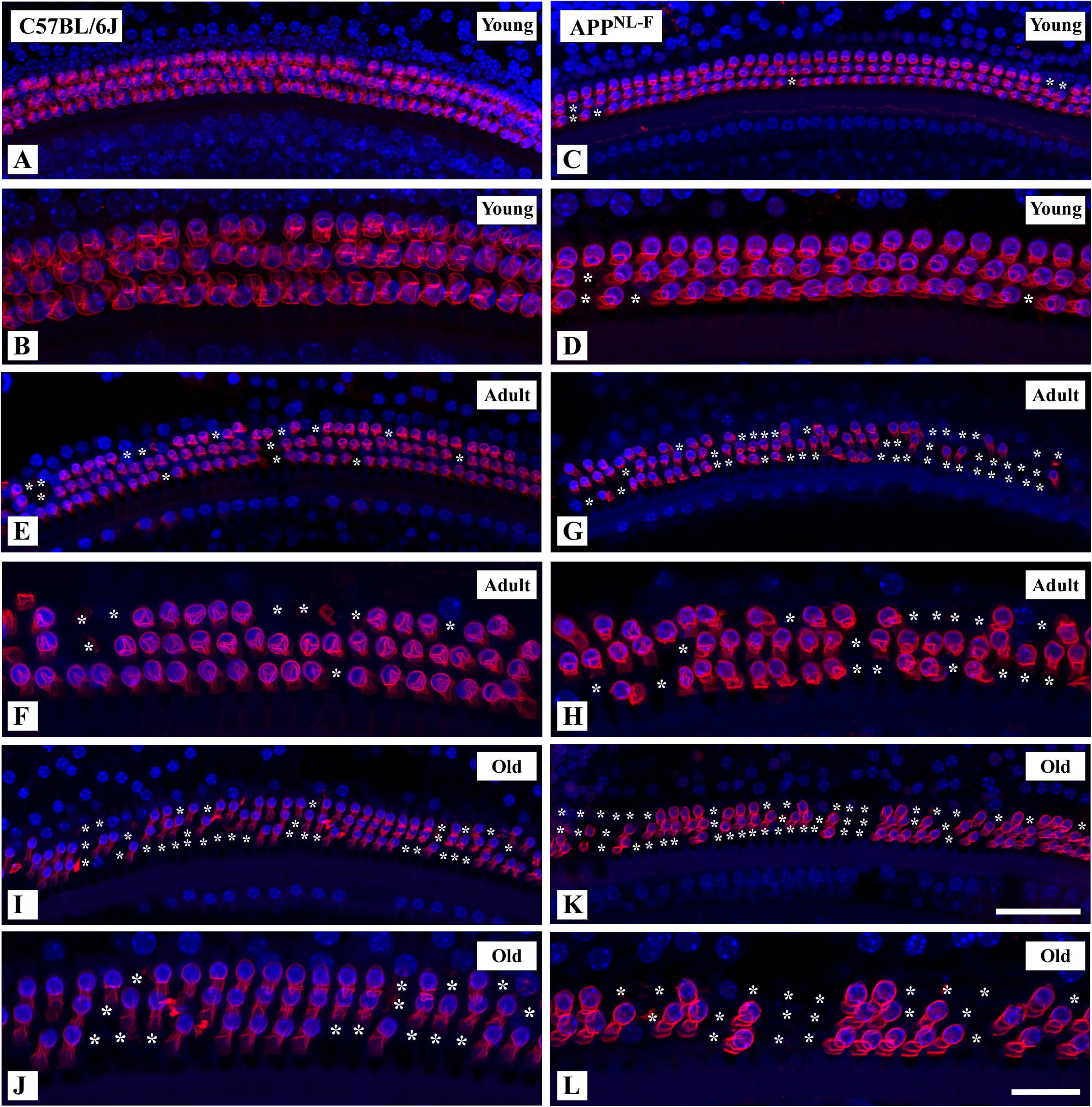
Whole-mount preparations of the organ of Corti. Confocal optical sections through a plane transversal to the sensory epithelium, at the upper level of the basal turn. Outer hair cell profiles immunostained with prestin (red magenta). Nuclei counterstained with DAPI (blue). A, B: Young C57BL/6J mouse. The three rows of outer hair cells are preserved (A). Prestin decorates plasma membranes (B). C, D: Young *App*^NL-F^ mouse. Outer hair cell architecture is overall well preserved (C), although outer hair cells are occasionally missing (D). E, F: Adult C57BL/6J mouse. Gaps (E) of one or two adjacent missing outer hair cells are seen (F). G, H: Adult *App*^NL-F^ mouse. Missing outer hair cells are more numerous (compare E and G). Long stretches with three or more adjacent missing outer hair cells are seen in the three rows (G, H). I, J: Old wild type C57BL/6J mouse. There is abundant outer hair cell loss (I). Long stretches of adjacent missing outer hair cells are frequently seen (J). K, L: Old *App*^NL-F^ mouse. Abundant outer hair cell loss is seen in the three rows (compare I, J and K, L). Scale bars: 50µm in K and 20 µm in L.

In young mice of both strains, the three rows of OHC rows were arranged in a continuous, homogeneous pattern from base to apex. Young *App*^NL-F^ mice occasionally showed small gaps of one or two missing OHCs in the basal and middle turns (Fig. 5A,B,G,H). In adult WT C57 mice, row continuity was disrupted by numerous gaps of one or two cells, affecting all three rows predominantly in the basal and middle turns (Fig. 5C,D). Adult *App*^NL-F^ mice showed markedly larger losses, with long stretches of consecutive missing cells across all rows (Fig. 5I,J). In old mice of both strains, widespread OHC loss was evident, with long discontinuous stretches separating clusters of remaining prestin-positive cells (Fig. 5E,F,K,L).

Quantification of OHC survival normalized to young animals is shown in Fig. 6 and Table 2. Survival was maximal in young mice of both strains, with no significant differences between them (Fig. 6A–C). In adult WT C57 mice, OHC survival was significantly reduced at the basal (85.41 ± 0.80 %) and middle (88.47 ± 0.88 %) turns, but not at the apex (92.08 ± 1.91 %) (Fig. 6; Table 2). In adult *App*^NL-F^ mice, the reduction in the basal turn was much larger (62.66 ± 6.47 %), significantly below both age-matched WT C57 mice and young mice of both strains (Fig. 6A; Table 2). Middle- and apical-turn survival (80.91 ± 1.05 % and 85.88 ± 3.19 %) was significantly reduced relative to young animals but did not differ from adult WT C57 mice.

**Figure 6.**
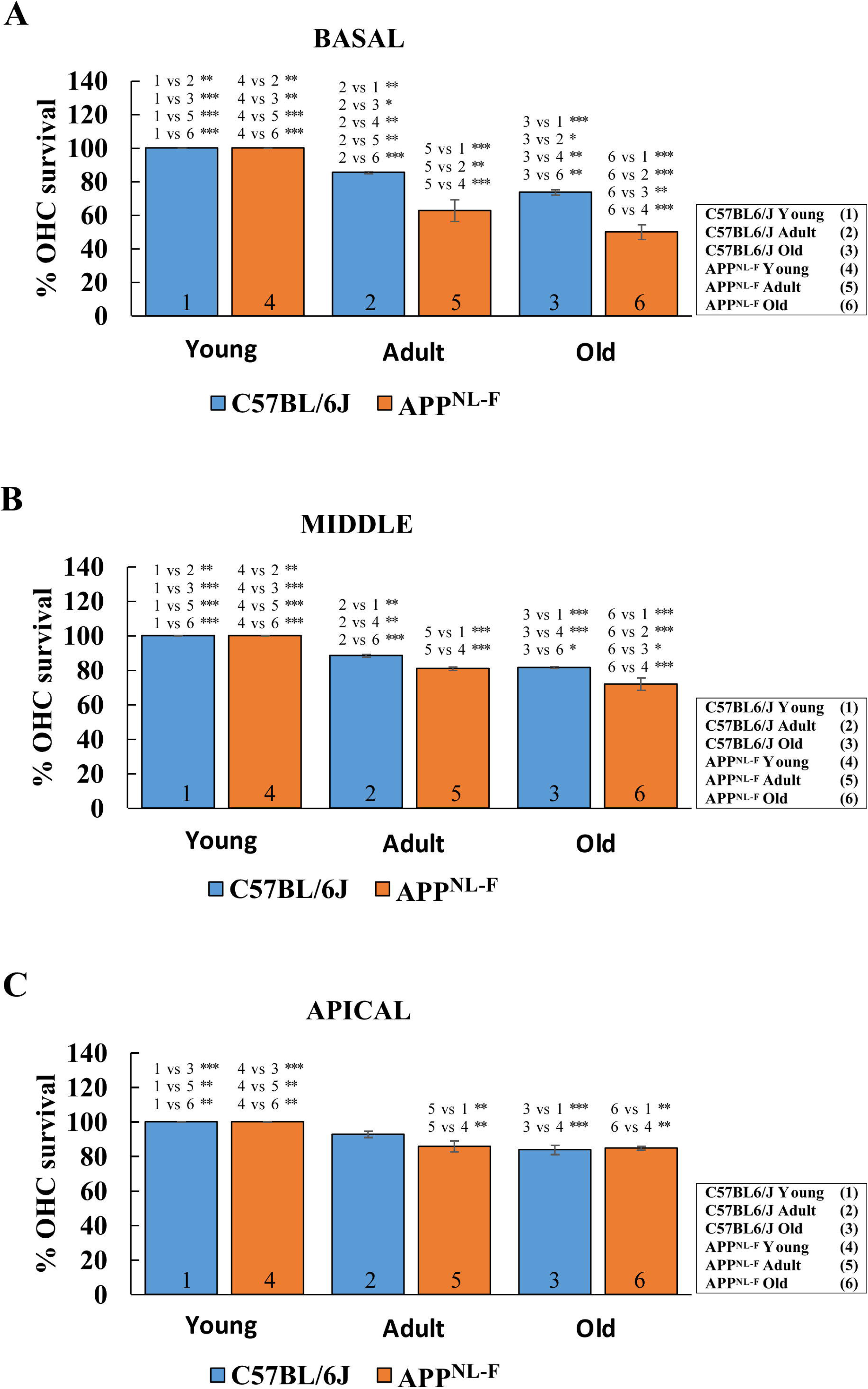
Outer hair cell (OHC) relative (%) survival in young, adult and old wild type C57BL/6J (blue) versus *App*^NL-F^ mice (orange) in the basal (A), middle (B) and apical (C) cochlear turns. A: Relative outer hair cell survival at the basal cochlear turn. Both mice strains show progressive significant reduction of outer hair cell survival from adult to old ages relative to the young age group. Progressive survival reduction with age in C57 BL/6J mice (blue bars) and in *App*^NL-F^ mice (orange bars) follows the same temporal pattern (1, 2 and 3 vs. 4, 5 and 6). However, percent outer hair cell survival values are significantly lower in adult and old *App*^NL-F^ mice (5, 6) than in age-matched wild type C57BL/6J (2,3). B: Relative outer hair cell survival at the middle cochlear turn. Overall, survival is higher than in the basal turn. However, there is still progressive reduction with age. In C57BL/6J mice (blue bars) survival is significantly reduced between young and adult (1, 2), but not between adult and old. In the middle cochlear turn of *App*^NL-F^ mice (orange bars), outer hair cell survival also is progressively reduced with age. At the adult age group, survival reduction does not differ significantly from C57BL/6J (2, 5). At the old *App*^NL-F^ group, however, outer hair cell survival in the middle turn is significantly lower than in old wild type C57BL/6J (3, 6). C: Relative outer hair cell survival at the apical cochlear turn. There is a small but significant reduction at the old age group both in C57BL/6J mice and *App*^NL-F^ mice (compare 3 with 1 and 6 with 4). At the apical turn, however, relative survival values do not differ significantly between both strains at the same age time points (compare 2, 5 vs. 3, 6). *p < 0,05; **p < 0,01; ***p < 0,001. See Table 2 for statistical tests details.

**Table 2.**
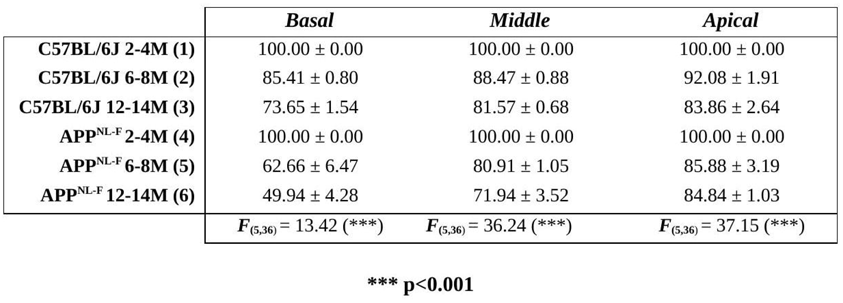
Mean ± SEM and ANOVA of the interaction between age and strain over the percentage of OHC survival relative to controls (young mice groups)

In old WT C57 mice, basal OHC survival further declined to 73.65 ± 1.54 %, significantly below values from both adult and young mice (Fig. 6A; Table 2); middle (81.57 ± 0.68 %) and apical turn (83.86 ± 2.64 %) survival were significantly lower than in young, but not than in adult, WT C57 mice (Fig. 6B,C). In old *App*^NL-F^ mice, basal-turn survival dropped to 49.94 ± 4.28 %, significantly below age-matched old WT C57 and younger animals of both strains, though not different from adult *App*^NL-F^. Middle-turn survival (71.94 ± 3.52 %) was significantly lower than in age-matched old WT C57 and in young and adult WT C57, but not than in adult *App*^NL-F^. Apical survival (84.84 ± 1.03 %) differed from young animals of both strains but not from adults of either genotype or from age-matched old WT C57 (Fig. 6; Table 2).

Overall, OHC loss progressed with age in both strains, with the greatest vulnerability at the basal turn. *App*^NL-F^ mice showed earlier and more pronounced OHC loss than age-matched WT C57 mice, consistent with accelerated cochlear sensory-cell degeneration in this strain.

### 3.4. Age-related cochlear microstructural changes in WT C57 and *App*^NL-F^ mice

H/E-stained modiolar sections (Figs. 7–9) revealed essentially identical cochlear morphology in young WT C57 and *App*^NL-F^ mice: the basal sensory epithelium showed the three OHC rows, the inner hair cell row, and the pillar cells flanking the tunnel of Corti; the SV and SL had normal cellular organization and density, the SV retaining its characteristic three-layered structure; and Rosenthal’s canal contained regularly arranged SG neuron somata of comparable morphology and packing density (Fig. 7).

**Figure 7.**
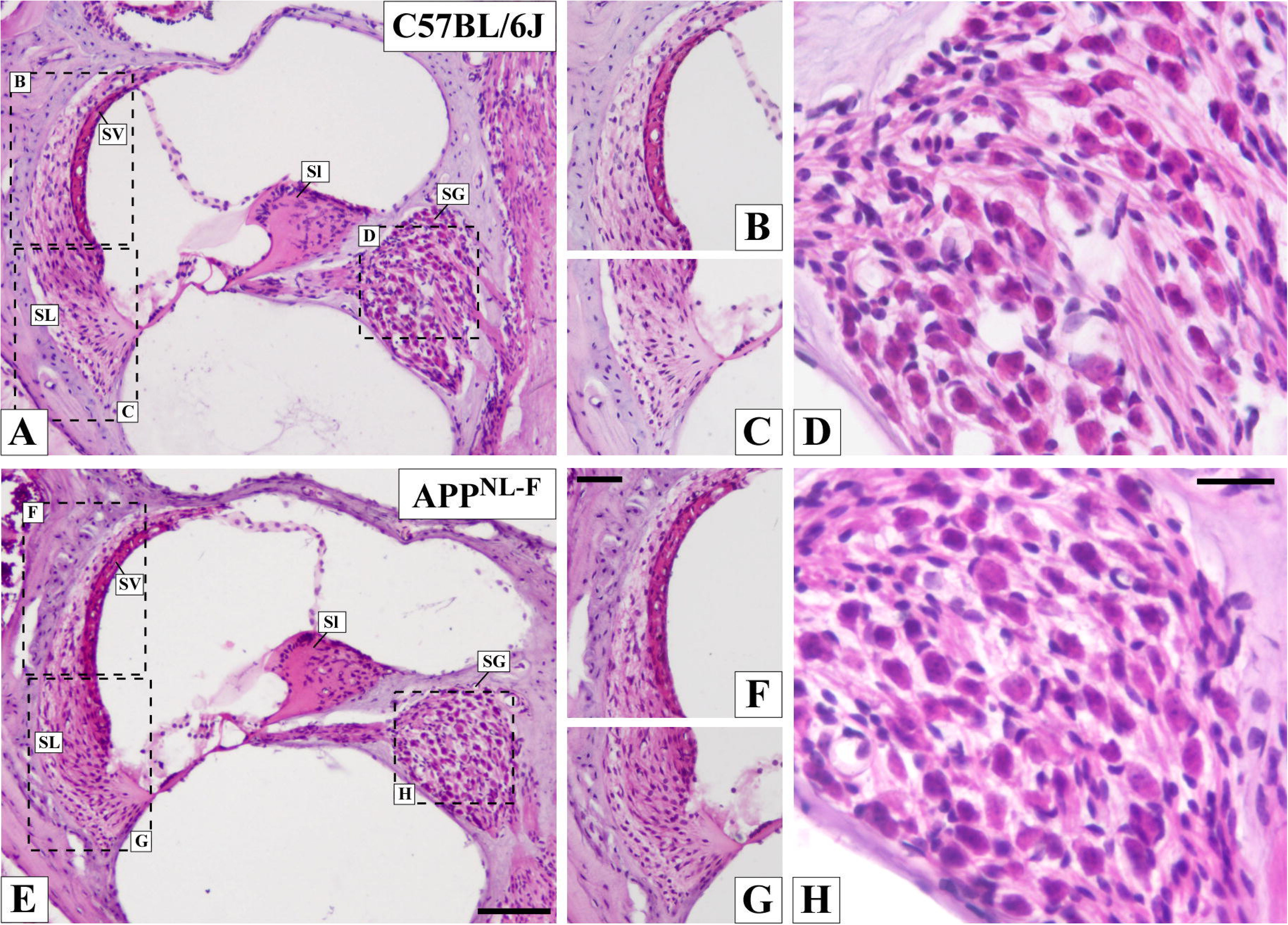
A, E: Cochlear mid-modiolar sections stained with hematoxylin-eosin showing cochlear histology in young C57BL/6J (A) and *App*^NL-F^ mice (E). The microarchitecture of the organ of Corti is well preserved in both strains. The SV in A and E has an identically regular cell arrangement (B, F inserts). Cell density and disposition in the SL in A and E are equally preserved (C, G inserts). In the SG in D and H, neuronal cell bodies packing and morphology are undistinguishable between both strains at the young age group. Abbreviations: SG, spiral ganglion; SL, spiral ligament; Sl: spiral limbus and SV; stria vascularis.

**Figure 8.**
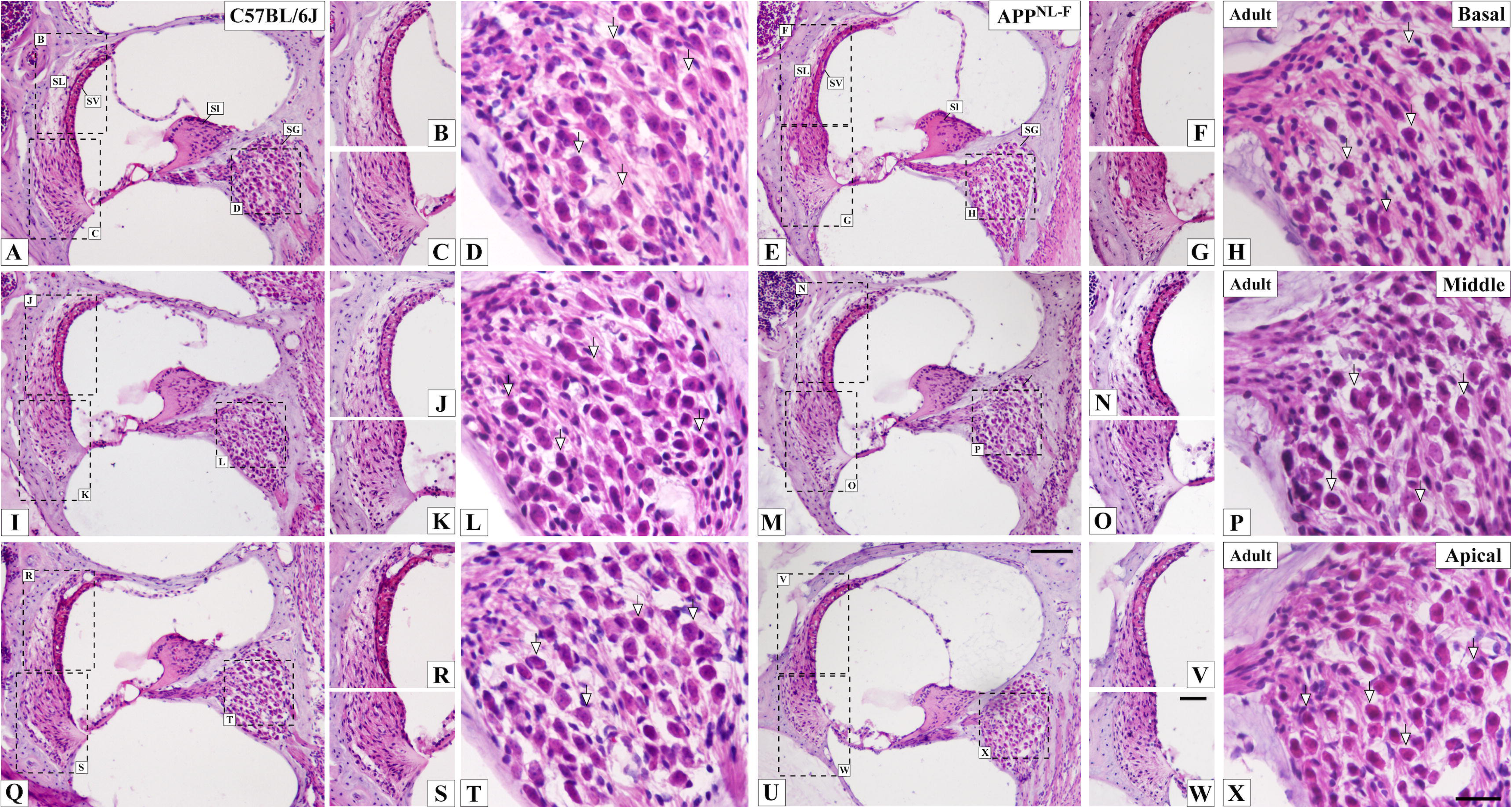
Cochlear para-modiolar sections stained with hematoxylin-eosin showing cochlear microstructure in adult C57BL/6J mice at the basal, middle and apical cochlear turns (A, I, Q) and in adult *App*^NL-F^ mice at the same levels (E, M, U). The organ of Corti architecture looks distorted in comparison with young mice, particularly at the basal turn (A, E). The SV and the SL do not show observable changes neither in C57BL/6J or in *App*^NL-F^ mice in any cochlear turn (compare inserts B, C with F, G; J, K with N, O and R, S with V, W). In the SG in A and E, neuron cell body morphology does not show differences between both strains at any cochlear turn (arrows in inserts D, L, T and H, P, X). Subtle changes in the packing of spiral ganglion neurons are observable at the cochlear base, with more visually empty extracellular space among them (D and H). Abbreviations: SG, spiral ganglion; SL, spiral ligament; Sl: spiral limbus and SV; stria vascularis. Scale bars: 100µm in U; 50µm in V and 25µm in X.

**Figure 9.**
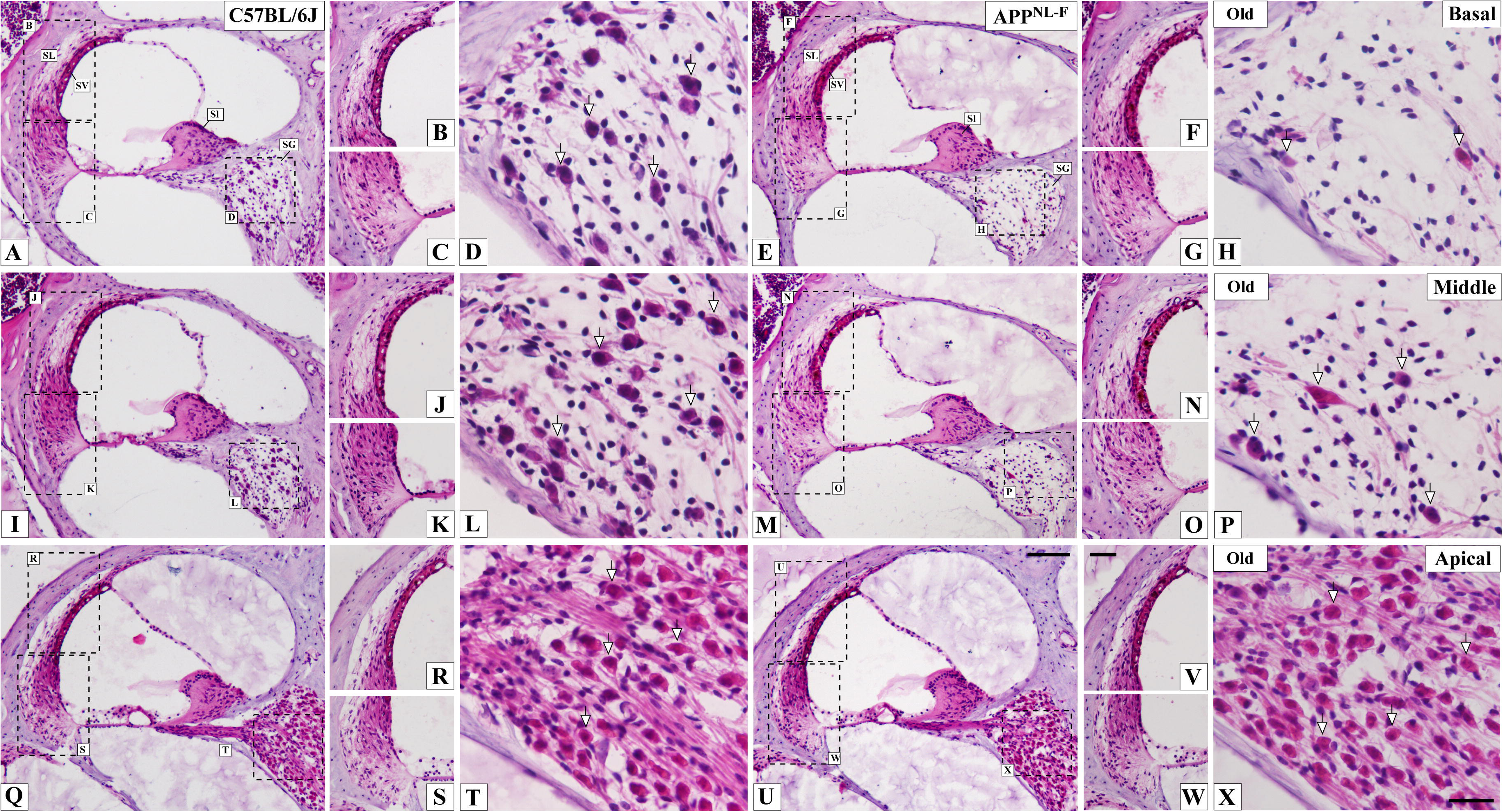
Cochlear para-modiolar sections stained with hematoxylin-eosin showing changes in cochlear microstructure in old C57BL/6J mice at the basal, middle and apical cochlear turns (A, I, Q) and in old *App*^NL-F^ mice at the same cochlear locations (E, M, U). The organ of Corti looks severely distorted at the basal and middle turns (A, E, I, M). In the basal and middle turns from the old *App*^NL-F^ mouse in the picture (E and M) only a flat epithelium remains. The SV (inserts B, J, R and F, N, V) shows more structural disorganization in the *App*^NL-F^ mouse, at the basal (F) and middle (N) turns, with packed variegated cell nuclei visible across its thickness (F, N). The SL does not show observable differences between old C57BL/6J and *App*^NL-F^ mice at any cochlear turn (compare inserts B,C and F, G; J,K and N,O; R, S and V, W). In the SG in A and E, large cellular depletion (arrows in inserts D, L, T) is seen in the C57BL/6J mouse at the basal (insert D) and middle (insert L) cochlear turns. In the SG at the apical turn, cellular depletion is less obvious (T). In the SG of the old *App*^NL-F^ mouse, cellular depletion at the base (H) and middle turn (P) looks larger, with fewer remaining cells (compare D, H and L, P). In the SG at the cochlear apex, cell packing density is slightly but still visibly diminished (compare T and X). Abbreviations: SG, spiral ganglion; SL, spiral ligament; Sl: spiral limbus and SV; stria vascularis. Scale bars: 100µm in U; 50µm in V and 25µm in X.

In adult mice, cochlear microstructure remained similar between strains and only slightly altered relative to young animals. Mild disorganization of the basal sensory epithelium was occasionally observed in both strains (Fig. 8A,E), while the SV, SL and Rosenthal’s canal appeared preserved at all cochlear levels (Fig. 8B–D, F–H, I, M, Q, T, U, X).

In old mice, both strains showed pronounced basal-turn degeneration, with frequent severe disruption of the sensory epithelium, occasional full denudation of the basilar membrane (more often in *App*^NL-F^), loss of the normal three-layered SV architecture, reduced SL cellularity — particularly in the type-IV fibrocyte region — and marked SG neuron depletion (Fig. 9A,B,D,E,F,H). These changes were consistently more severe in *App*^NL-F^ mice, in which irregular clusters of nuclei within the intermediate cell layer of the SV were frequent (Fig. 9F) and near-complete SG neuron depletion in the basal turn was not uncommon (Fig. 9D,H). Similar but somewhat less severe alterations were present at the middle turn (Fig. 9I–P), whereas the apical sensory epithelium, lateral wall and SG remained the best preserved in both strains (Fig. 9Q–X). Thus, age-related structural degeneration of the sensory epithelium, lateral wall and spiral ganglion occurred in both strains, but was more severe — especially at the basal turn and in the spiral ganglion — in *App*^NL-F^ mice.

### 3.5. Comparative spiral ganglion neuron survival in WT C57 and *App*^NL-F^ mice

In young mice, SG neuron profile counts in H/E-stained modiolar sections did not differ between WT C57 and *App*^NL-F^ mice and served as the reference (100 %) for normalization at each cochlear turn (Fig. 10A–C; Table 3). In adult mice, normalized SG survival did not differ significantly from the young reference in either strain, nor between adult WT C57 and adult *App*^NL-F^ mice, at any cochlear turn (Fig. 10A–C; Table 3).

**Figure 10.**
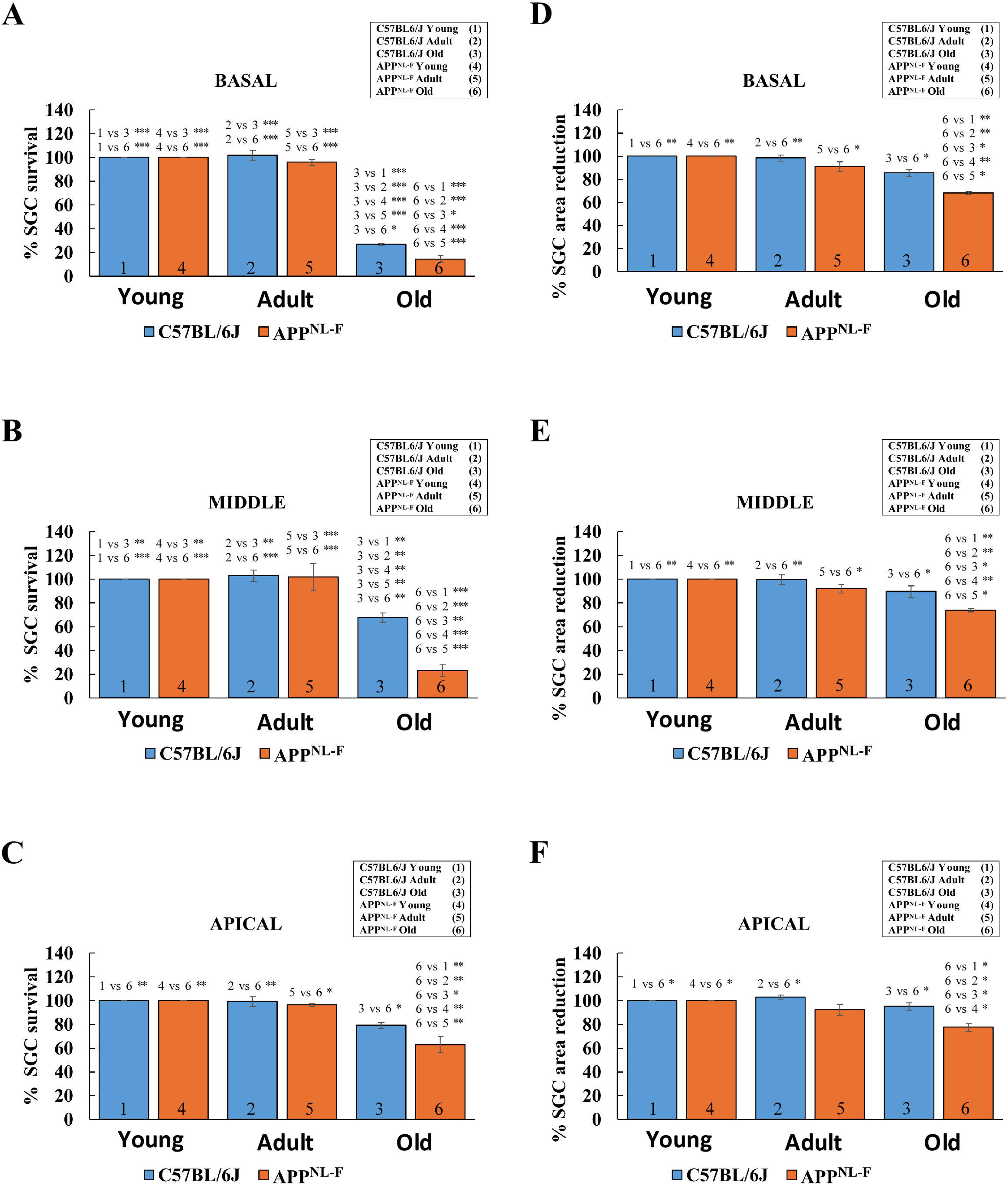
Relative (%), spiral ganglion cell (SGC) survival at basal (A) middle (B) and apical cochlear (C) locations and relative (%), spiral ganglion cell (SGC) area changes (D, E, F) in young, adult and old C57BL/6J (blue) and *App*^NL-F^ mice (orange). In old mice of both strains, there is large, significantly diminished SGC survival at the cochlear base relative to adult and young mice (A: 3, 6 vs. 2, 5 and 1, 4). SGC relative survival is significantly lower in old *App*^NL-F^ (A, 6) than in C57BL/6J mice (A, 3). In middle cochlear turns, the timeline pattern of SGC survival is similar (B: 3, 6 vs. 2, 5 and 1, 4). SGC relative survival is significantly lower in old *App*^NL-F^ (B, 6). Percent SGC survival values, however, are larger than at the cochlear base. At the cochlear apex (C), relative SGC survival is close to maximum in young, adult and old wild type C57BL/6J mice (C: 1, 2, 3). A small percent reduction in old mice of this strain, is not significant relative to adult and young (C, 3). At the cochlear apex of old *App* ^NL-F^ mice, however, there is significant reduction of SGC survival (C: 6 and 3). SGC area reduction (D, E, F) is significant in old *App*^NL-F^ mice at the base, middle and apical portions of the cochlea (D, E, F) relative to old C57BL/6J mice (D, E, F: 3, 6) ant to adult and young mice of both strains. *p < 0,05; **p < 0,01; ***p < 0,001. See Table 3 and Table 4 for statistical tests details.

**Table 3.**
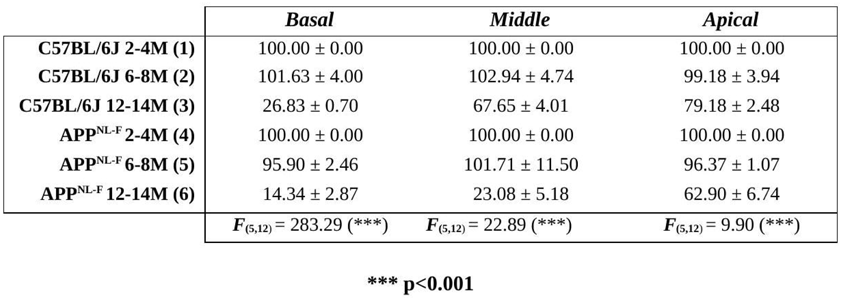
Mean ± SE and ANOVA of the interaction between age and strain over the percentage of SG neuron survival relative to controls (young mice groups).

**Table 4.**
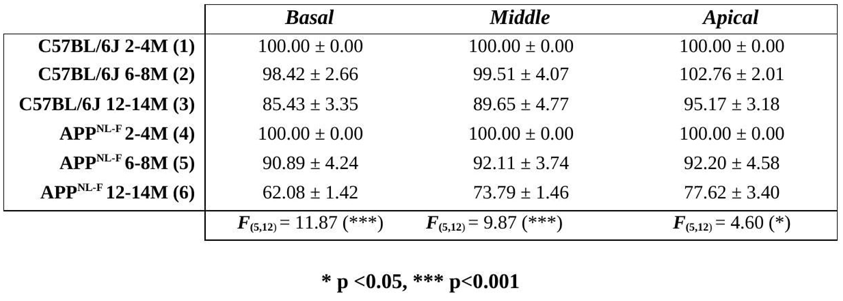
Mean ± SE and ANOVA of the interaction between age and strain over the percentage of SG neuron area reduction relative to controls (young mice groups).

In old mice, SG neuron survival declined markedly in both strains along the full length of Rosenthal’s canal. In old WT C57 mice, survival was lowest at the basal turn (26.83 ± 0.70 %), significantly reduced relative to young and adult animals of both strains (Fig. 10A). In old *App*^NL-F^ mice, basal-turn survival fell further to 14.34 ± 2.87 %, significantly lower than in age-matched WT C57 and in young and adult animals of both strains (Fig. 10A; Table 3). Middle-turn survival in old WT C57 averaged 67.65 ± 4.01 %, significantly below young and adult values; in old *App*^NL-F^ mice the decrease was much greater (23.08 ± 5.18 %) and significantly larger than in age-matched WT C57 (Fig. 10B; Table 3). At the apex, survival in old WT C57 remained relatively preserved (79.18 ± 2.48 %, not significantly different from young or adult groups), whereas in old *App*^NL-F^ mice it dropped significantly (62.90 ± 6.74 %), both relative to age-matched WT C57 and to young and adult animals of both strains (Fig. 10C; Table 3).

SG neuron loss therefore progressed with age in both mice strains along the base-to-apex gradient typical of ARHL but was substantially greater in *App*^NL-F^ mice at every cochlear level, pointing to increased auditory-nerve vulnerability to both age- and Alzheimer-related neurodegenerative mechanisms in this strain.

### 3.6. Changes in spiral ganglion neuron cell body size in WT C57 and *App*^NL-F^ mice

SG neuron cross-sectional areas in young mice were uniform across cochlear turns and did not differ between strains, providing the reference for normalization of age-related changes (Fig. 10D–F; Table 4). In adult mice, SG soma areas remained close to young reference values in both strains, with no significant differences at any cochlear turn (Fig. 10D–F).

In old WT C57 mice, the reduction in SG soma area relative to young animals did not reach significance at any cochlear turn (Fig. 10D–F; Table 4). In contrast, old *App*^NL-F^ mice showed significant reductions at the basal, middle and apical turns relative to age-matched old WT C57 and to young and adult animals of both strains, with the sole exception of the comparison with adult *App*^NL-F^ at the apex (Fig. 10D–F; Table 4).

Aging therefore had little effect on the soma size of surviving SG neurons in WT C57 mice. However, neuronal atrophy awas significant across the cochlear spiral in old *App*^NL-F^ mice — a degenerative feature accompanying the marked SG neuron loss observed in this Alzheimer-disease model.

### 3.7. Accelerated stria vascularis thinning in *App*^NL-F^ mice

SV thickness measurements for all turns, ages and strains are summarized in Table 5 and Fig. 11. In young animals, mean SV thickness was ∼22–24 µm across cochlear turns in both strains, with no significant differences (Fig. 11A–C; Table 5).

**Figure 11.**
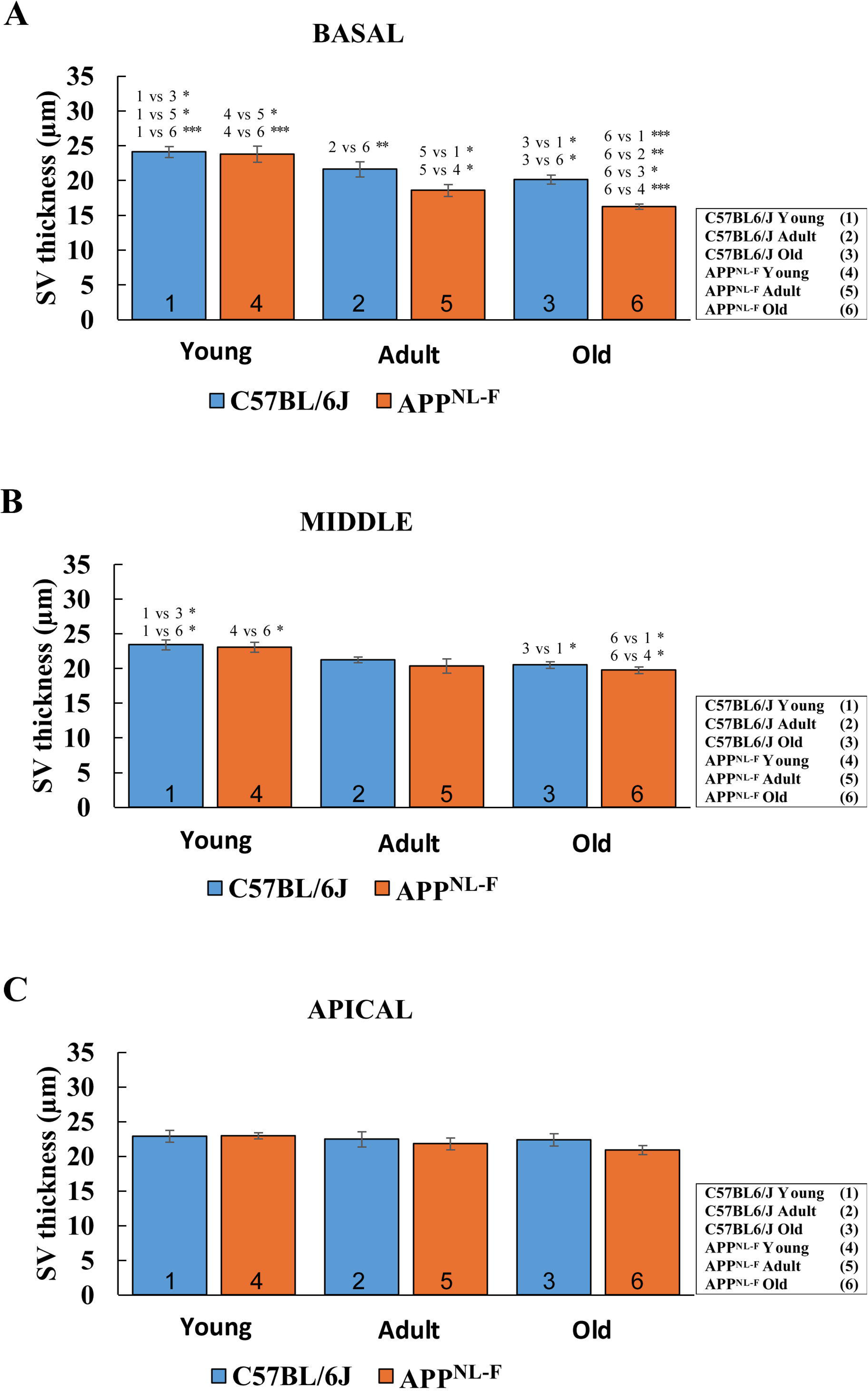
Thickness changes in the stria vascularis (SV) at basal (A) middle (B) and apical cochlear locations (C) in young, adult and old C57BL/6J (blue) and *App*^NL-F^ mice (orange). At the cochlear baseof C57BL/6J mice, the SV is significantly reduced in thickness in old relative to young mice (A: 3, 1). In *^A^pp* N^L-F^ mice SV thickness reduction at the cochlear base is significant at the adult group relative to young mice (A: 5, 1,4). In old mice of this strain, thickness reduction of the SV at the cochlear base is significantly larger than in old C57BL/6J mice. In the SV at middle cochlear locations, there is a slight but significant thickness reduction at old ages in both mice strains (B: 3, 6, 1, 4). At the apical turn (C), SV thickness does not change significantly in either strain during aging. See Table 5 for statistical tests details.

**Table 5.**
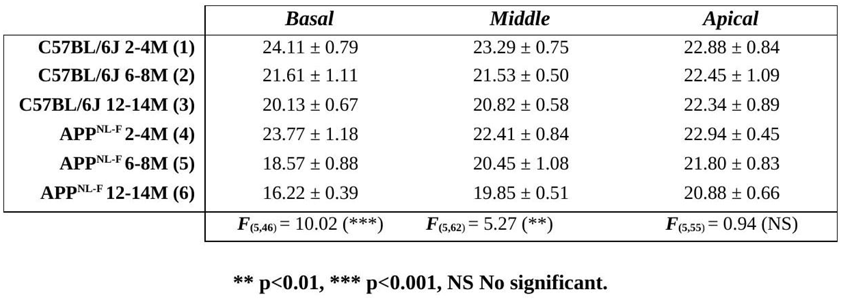
Mean ± SE and ANOVA of the interaction between age and strain over stria vascularis thickness (μm).

In adult WT C57 mice, SV thickness remained close to young values at all turns (Fig. 11A–C). Adult *App*^NL-F^ mice, by contrast, showed a significant reduction at the basal turn (18.57 ± 0.88 µm) relative to young animals of both strains; the comparison with age-matched adult WT C57 did not reach significance. Middle- and apical-turn SV thickness did not differ between adult groups or from young animals (Fig. 11A–C; Table 5).

In old mice, basal-turn SV thickness declined to 20.13 ± 0.67 µm in WT C57 and to 16.22 ± 0.39 µm in *App*^NL-F^ mice, the latter being significantly reduced relative both to age-matched WT C57 and to young animals of both strains (Fig. 11A). In the middle turn, both old WT C57 and old *App*^NL-F^ mice (19.85 ± 0.51 µm) showed significant thinning relative to young animals (Fig. 11B). No significant differences were detected at the apex in any age or strain group (Fig. 11C).

Thus, *App*^NL-F^ mice show earlier and more pronounced SV atrophy, particularly at the basal turn, indicating increased vulnerability of the cochlear lateral wall in this Alzheimer model that may contribute to the functional hearing deficits observed with aging.

### 3.8. Vascular alterations in the stria vascularis in WT C57 and *App*^NL-F^ mice

Because the basal SV showed the greatest age-related thinning (section 3.7) and paramodiolar sections from old animals frequently revealed enlarged capillary lumina (Fig. 11), we measured SV capillary lumen area in the three cochlear turns on phalloidin/DAPI-stained sections (Fig. 12; Table 6).

**Figure 12.**
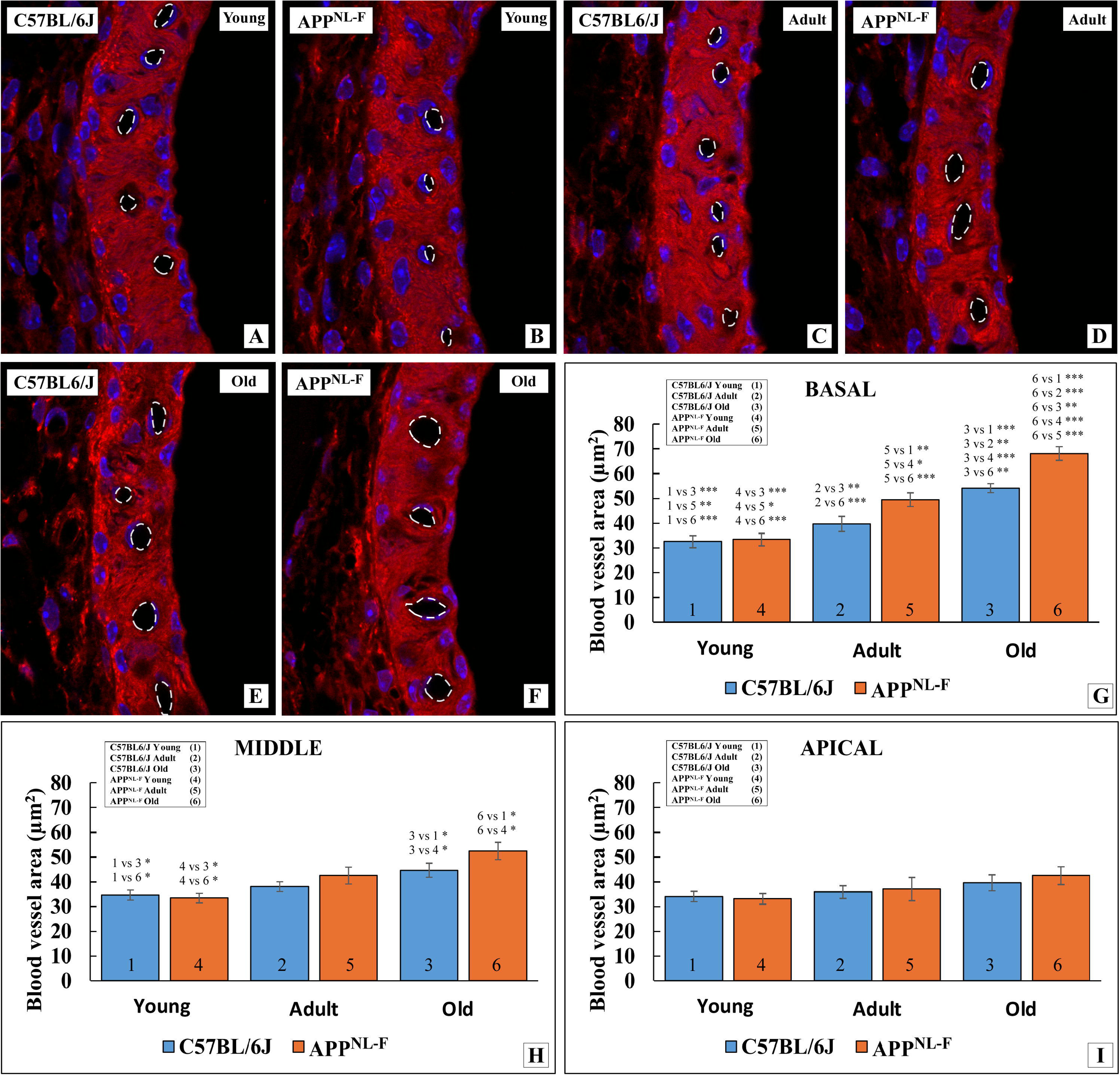
Sections through the stria vascularis of C57BL/6J and *App*^NL-F^ mice at the cochlear base at young (A-B), adult (C-D) and old (E-F) ages. Fluorescent phalloidin staining (red) and nuclear counterstaining with DAPI (blue). Laser scanning confocal microscopy imaging. Dashed white irregular circles mark capillary lumen limits, drawn at the border between the black, empty lumen space and the immediately adjacent red (phalloidin) or blue (nuclei) staining, taken respectively as the luminal face and nuclei of the capillary endothelial cell. Capillary lumen size looks larger with age (A, C, E and B, D, F). G-I: Bar graphs showing capillary blood vessel area during aging in basal, middle and apical portions of the stria vascularis in C57BL/6J (blue) and in *App*^NL-F^ mice (orange). In the basal turn (G), average capillary blood vessel area increases progressively with age both in wild type C57BL/6J (G: 1, 2^, 3)^ and in *App*^NL-F^ mice (G: 4, 5, 6). At the old age group, capillary blood vessel area is s^igni^ficantly larger in *App*^NL-F^ mice. In middle portions of the stria vascularis (H) capillary lumen area becomes significantly larger only at old ages (H: 5, 6 and 1, 2), with no^dif^ ferences between C57BL/6J and *App*^NL-F^ (H: 5, 6). At the cochlear apex (I), capillary blood vessel area does not change significantly between both mice strains at any age group. *p < 0,05; **p < 0,01; ***p < 0,001. See Table 6 for details on statistical tests.

In young mice, SV capillaries displayed narrow, oval-to-round lumina outlined by phalloidin-labelled cell contours and DAPI-stained endothelial nuclei (Fig. 12A,B). Lumen area was highly similar between strains at every turn (∼33–35 µm²) (Fig. 12G–I; Table 6). In adult mice, lumen morphology remained broadly similar, with occasional mildly enlarged profiles in both strains (Fig. 12C,D). Adult WT C57 lumina were slightly but non-significantly larger than in young animals (basal 39.69 ± 3.02 µm²; middle 38.05 ± 2.03 µm²; apical 35.90 ± 2.58 µm²). In adult *App*^NL-F^ mice, basal-turn lumen area rose significantly to 49.44 ± 2.76 µm² relative to young animals, although the comparison with age-matched adult WT C57 did not reach significance; middle- and apical-turn lumen areas (42.52 ± 3.34 µm² and 37.06 ± 4.70 µm²) did not differ from adult WT C57 or young groups (Fig. 12G–I; Table 6).

**Table 6.**
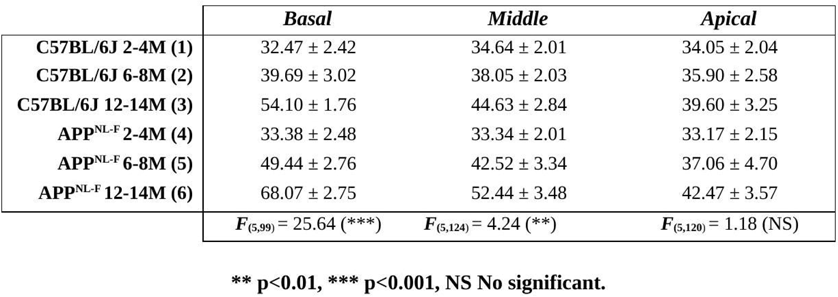
Mean ± SE and ANOVA of the interaction between age and strain over stria vascularis blood vessel area (μm^2^).

In old animals, SV capillary lumina were markedly enlarged (Fig. 12E,F), more so in *App*^NL-F^ mice. Basal-turn lumen area rose to 54.10 ± 1.76 µm² in old WT C57 (significantly greater than in adult WT C57 and in young animals of both strains) and to 68.07 ± 2.75 µm² in old *App*^NL-F^, significantly exceeding age-matched WT C57 and both adult and young animals (Fig. 12G; Table 6). In the middle turn, both old groups (WT C57: 44.63 ± 2.84 µm²; *App*NL-F: 52.44 ± 3.48 µm²) were significantly above young animals but did not differ from adult animals or from each other (Fig. 12H). At the apex, lumen area showed only a non-significant increase in old *App*^NL-F^ mice (Fig. 12I; Table 6).

SV capillary lumen area therefore enlarged progressively with age, preferentially at the basal and middle turns, and was significantly greater in *App*^NL-F^ mice at the basal turn. This vascular remodeling may reflect impaired SV homeostasis and could contribute to the accelerated auditory decline observed in *App*^NL-F^ mice.

## 4. Discussion

We provided evidence that brain Aβ pathology linked to AD accelerates ARHL traits in the auditory receptor organ. An initial consideration is the novelty and robustness of the animal model utilized. The “knock in” *App*^NL-F^ mouse was generated to reproduce Aβ pathology in regions of the brain cortex homologous to those primarily involved in AD in humans [25,29–31]. Aβ pathology was primed by “knocking in” at its specific locus, through homologous recombination, a “humanized” *App* gene containing mutations in the Aβ region of the sequence [31]. The resulting APP product is prone to dysregulated proteolytic attack, with pathologic accumulation of Aβ [30,31]. Because transcription is governed by the mouse promoter at its specific chromosomal loci, there is no APP overexpression. Thus, abnormal Aβ processing and deposition takes place mostly in brain regions involved in AD, in which neurons express “physiological” levels of APP [25,29–31]. It has been proposed that the *App*^NL-F^ mouse reproduces early stages of AD [25,29]. Like other genetically modified mice strains, the *App*^NL-F^ mouse was generated from the “wild type” genetic background of the C57BL/6J mouse [31]. The availability and well-known biology of this inbred mouse strain [43,44] makes it convenient for this aim [28]. It is known that the C57BL/6J mouse develops early progressive hearing loss [32], which is fully established between six and eight months [45,46]. Hearing loss originates, at least in part, from a spontaneous, carried-over mutation in the cadherin 23 gene (*Cdh23*) [46], which impairs mechano-electrical transduction in auditory hair cells. Auditory threshold elevations start in the high frequency range, progressing to intermediate and lower frequencies [35,47]. There is progressive auditory hair cell and spiral ganglion neuron loss spreading from the cochlear base to the apex [47]. All these are traits of peripheral human ARHL, of which the C57BL/6J mouse is regarded as a reliable model [45]. We reasoned that convergence in the *App*^NL-F^ mouse of induced AD Aβ pathology and traits of ARHL from the “wild type” C57BL/6J background (WT C57) may constitute a model of AD/ARHL “comorbidity”, allowing to dissect out possible interactions between both. This is relevant in view of recent epidemiological evidence supporting that hearing loss in middle stages of life in humans is the main modifiable risk factor for dementia in midlife, including AD [20]. Once characterized, such animal model may become a valuable tool to unravel whether and how ARHL and AD influence each other, a fundamental step for preventative and therapeutic strategies to handle cognitive impairment [4,15,20,48].

Within this conceptual framework, we gathered evidence showing that: a) loss of hearing sensitivity, i.e. increased auditory thresholds and large threshold shifts in adult WT C57 mice, is accelerated in age-matched *App*^NL-F^ mice; b) there is increased loss of OHCs and SG neurons, as well as damage to the SV in adult and old *App*^NL-F^ mouse, which may explain accelerated hearing loss in the *App*^NL-F^ strain. As no interfering phenotypes have been reported in this *App* “knock in” mouse, we propose that brain AD Aβ pathology may trigger, so far unknown, pathophysiological events “backpropagated” to the auditory receptor, leading to accelerated ARHL. This, in turn, may further accelerate cognitive impairment. Implications of such potential “vicious circle”, emerging from results in this “dual” AD/ARHL animal model, will be further discussed.

### 4.1. Auditory threshold elevation during aging is accelerated in *App*^NL-F^ mice

Absolute threshold and threshold shift levels showed that hearing sensitivity was equally preserved at young ages in WT C57 and *App*^NL-F^ mice. Adult WT C57 mice, i.e., around seven months of age, had significantly elevated thresholds to all tested tonal frequencies, relative to young WT C57 mice. Overall, this supports previously reported early ARHL-like attenuation of hearing sensitivity in WT C57 mice [36]. On the other hand, adult *App*^NL-F^ mice, also around seven months of age, had thresholds both to clicks and tonal frequencies, significantly more elevated than their young *App*^NL-F^ and WT C57 mice counterparts. Importantly, at this adult age, auditory thresholds were significantly higher in *App*^NL-F^ than in WT C57 mice. Also, threshold shifts for clicks and individual frequencies were considerably larger in adult *App*^NL-F^ than in age-matched WT C57 mice. Thus, early ARHL-like degradation of hearing sensitivity in adult WT C57 mice is further accelerated in age-matched *App*^NL-F^. As mentioned, because there are no known confounding phenotypic traits in the *App*^NL-F^ strain [29,31], accelerated loss of hearing sensitivity in adult *App*^NL-F^ is most probably linked to central Aβ pathology.

This notion gains additional support when hearing thresholds in old mice are considered. In old WT C57 mice, around 13.4 months of age, average thresholds increased significantly relative to adults, supporting hearing degradation evolving from adulthood throughout the aging process in this mouse strain [45,46]. Old *App*^NL-F^ mice, aged around 14 months, also showed significant elevations in auditory thresholds relative to adult WT C57. However, they did not differ significantly from those in adult *App*^NL-F^ mice. Likewise, threshold elevations were not significantly different between old *App*^NL-F^ and old WT C57 mice. In addition, differences in auditory threshold shifts were, overall, larger between young and adult *App*^NL-F^ than between adult and old *App*^NL-F^. Also, correlation coefficients between age and threshold increases showed lower values for *App*^NL-F^ mice than for WT C57, particularly at the highest tested frequencies of 8 and 16 kHz. This may be an indication that a steep, linear loss of hearing sensitivity is reached earlier in *App*^NL-F^ mice, particularly at higher frequencies. Therefore, ARHL is accelerated in *App*^NL-F^ mice, again most likely because of Aβ pathology, so that diminished hearing sensitivity equivalent to old WT C57 appears earlier in *App*^NL-F^ mice, at the adult age of around 7.3 months. Beyond this timepoint, the rate of hearing loss in old *App*^NL-F^ slows down, at least until the selected timepoint of 14 months.

Reduced hearing sensitivity has been previously reported in some strains of AD transgenic mice with induced overexpression of mutated APP [39], which, as previously mentioned, leads to extensive deposits of Aβ throughout the central nervous system. Transgenic mice provide valuable information about pathological roles of Aβ However, APP overexpression may introduce confounding factors as far as Aβ pathology in AD is concerned, because there is also Aβ deposition in brain regions which are not originally involved in AD [25,29]. As mentioned, in the *App*^NL-F^ mouse and related “knock in” strains, Aβ accumulation is largely restricted to regions of the mouse brain cortex homologous to those involved in AD in humans. Therefore, acceleration of ARHL throughout aging in *App*^NL-F^ mice reported in this study may be attributable to Aβ pathology directly linked to AD, particularly at early stages [29] and not just to widespread deposition of Aβ, including auditory pathways, like in “standard” transgenic mice. In the related “knock in” *App*^NL-G-F^ mouse [25], high frequency auditory thresholds are increased relative to control C57BL/6J mice past six months of age [39]. Interestingly, changes in protein profile content in the perilymph, relative to WT C57 controls also have been found [38]. For instance, the concentration of proteins related to cochlear inflammation was higher in *App*^NL-G-F^ mice. This suggests further exacerbation of aging events in the cochlea [3] by the central Aβ pathology of AD. However, mechanisms through which this AD proteinopathy may affect auditory aging remain to be explored.

### 4.2. Increased outer hair cell loss and spiral ganglion neuron loss during aging in *App*^NL-F^ mice matches reduced hearing sensitivity

In adult WT C57 mice, OHC survival was significantly reduced. OHC loss was concentrated at the basal turn and to a lesser extent at the middle turn, with OHC survival around 85% and 88% respectively, relative to young WT C57 mice. OHC loss at the apical turn was not significant. This basal-to-apical pattern of OHC loss progressed further through aging, with survival rates in old WT C57 mice significantly reduced to less than 74% at the basal turn, below 82% at the middle turn, and 84% at the apical turn, relative to young WT C57 mice. Previous reports confirm a progressive, age-linked basal-to-apical gradient of OHC loss, starting early in adulthood in the WT C57 mouse [34,47], matching progressive increase of absolute thresholds and larger threshold shifts. Early, progressive ARHL-like OHC loss, characteristic of the WT C57 mouse was dramatically increased and accelerated in the *App*^NL-F^ mouse. OHC survival was lowest at the basal turn, with rates below 63%. At the middle turn, OHC survival in *App*^NL-F^ mice was below 81% and at the apical turn it was around 86%. Thus, OHC survival was significantly diminished throughout the cochlea in adult *App*^NL-F^ mice relative to adult WT C57, with OHC survival reduction encroaching the apical turn. Of note, OHC survival reduction in adult *App*^NL-F^ mice resembles that in old WT C57 mice, supporting acceleration of ARHL. In old *App* ^NL-F^ mice, OHC survival at the basal turn was below 50%, much lower than in age-matched old WT C57 mice. At the middle turn, OHC survival also was lower than in old WT C57, whereas at the apical turn it did not differ from old WT C57 mice. Therefore, loss of OHCs in ARHL is further accelerated and potentiated by co-existing AD Aβ pathology. It remains to be seen how aging-related mechanism of auditory hair cell damage and loss [49] may be potentiated by central Aβ pathology.

Another landmark of ARHL is damage and loss of SG neurons [3,49,50]. We found that relative SG neuron survival did not differ between adult and young WT C57 mice in any Rosenthaĺs canal segment, from base to apex. However, there was a dramatic decrease in relative survival rate of SG neurons between adult and old WT C57 mice, with a clearcut basal-to-apical pattern. Relative SG neuron survival in old WT mice was barely 27% at the cochlear base, slightly above 68% at the middle portion and 79% at the apical portion. Progressive loss of SG neurons during aging in WT C57 has been reported previously [50,51]. Our results support that SG neuron loss linked to aging is out of register with OHC loss [49–52]. The former manifests later and more abruptly, between the adult and old age stages [50]. Past a developmental time window, many SG neurons survive in the absence of sensory cells during adulthood and this may also apply to aging [49,51,52].

In *App*^NL-F^ mice, SG neuron loss followed a temporal pattern like that in WT C57 mice. However, old *App*^NL-F^ mice had significantly lower SG neuron survival rates. In basal portions of the SG, survival was slightly above 14%. In middle and apical portions, it was around 23% and 63% respectively. It is noticeable that in old *App*^NL-F^ mice, reduced neuron survival in the middle cochlear portions of Rosenthaĺs canal approximates that at the cochlear base in adult WT C57 mice. Thus, there was still a basal-to-apical SG neuron survival gradient in old *App*^NL-F^ mice, but relative survival was lower throughout the SG than in WT C57 mouse. Therefore, the Aβ proteinopathy may reduce SG neuron survival during aging. This further constrains input to central auditory nuclei, which may contribute to worsen central auditory and cognitive processing [22].

The trophic status of surviving SG neurons was tested using relative SG neuron cell body area as a proxy. In old WT C57 mice there was a slight, statistically not significant reduction of SG neuron cell body area relative to adult and young mice. One interpretation is that SG neurons surviving the natural aging process may be relatively more resilient to atrophy and thus more likely to maintain functional capabilities for a longer time. In contrast, in old *App*^NL-F^ mice there was a significant reduction in SG neuron cell body area throughout the SG, relative to old WT C57 mice. Insofar reduced cell body area may be interpreted as increased SG neuron atrophy in old *App*^NL-F^ mice, we hypothesize that central Aβ pathology may further limit functional capabilities of surviving SG neurons through yet unknown routes.

### 4.3. Increased atrophy in the stria vascularis of *App*^NL-F^ mice matches accelerated hearing loss

In WT C57 mice, degeneration and atrophy of the SV was histologically detectable only at the old age group. In old WT C57 mice, the SV showed structural disorganization of its cell layers and significantly diminished thickness, especially at its basal portion. At the mid-cochlear portion, the SV was slightly but still significantly reduced in thickness, whereas in the apical turn thickness did not change. Reduced SV thickness in old WT C57 mice has been reported by others [53–56], supporting that in this mouse model, aging-related gross structural damage to the SV is considerably delayed relative to OHC damage and loss. Thickness has been challenged as a predictor of the actual functional status of the SV [54,55,57]. However, even if aging interferes with the generation of the endocochlear potential before any histological changes in thickness are detected in the SV, it remains a useful indicator of the timeline of peripheral ARHL [58], like in the WT C57 mouse model.

In *App*^NL-F^ mice, degeneration and atrophy of the SV were accelerated and more intense. Different to WT C57, there was already a reduction in the thickness of the SV at the basal turn in adult *App*^NL-F^ mice, relative to young mice. Such diminished average SV thickness in adult *App*^NL-F^ mice did not differ significantly from that in old WT C57 or *App*^NL-F^ mice. On the other hand, in old *App*^NL-F^ mice SV thickness was further reduced relative to age-matched old WT C57 mice, at least at the cochlear base. Also, there was a slight reduction in the thickness of the SV at the middle turn, rather like that found in WT C57 mice, whereas no differences in thickness were seen at the apical turn, relative to mice of both strains at the adult or young age groups. Thus, the intensity of SV atrophy in old WT C57 mice determined by thickness measurements, is attained considerably earlier, at adult stages in *App*^NL-F^ mice and such SV atrophy is larger in old *App*^NL-F^ mice than in old WT C57. Therefore, aging-linked damage to the SV also seems to be accelerated and intensified by AD central Aβ pathology.

### 4.4. Capillaries in the stria vascularis in aged *App*^NL-F^ mice vs. WT C57 mice

Observable differences in the lumen size of capillary blood vessels integrated in the structure of the SV, prompted us to measure changes in lumen surface area. In young mice, capillary lumen area did not differ between WT C57 and *App*^NL-F^ strains. Average lumen size was very similar in basal, middle and apical portions of the SV. In WT C57 mice, measured SV capillary lumen area increased with age in basal and middle portions. In old WT C57 mice, lumen size was larger than in young and adult mice, which did not differ among them, both in basal and middle portions of the SV. In *App*^NL-F^ mice, lumen size increased significantly between young and adult mice and between these and old mice. Lumen size was larger in old *App*^NL-F^, with significant differences with age-matched old WT C57 mice. SV capillary degeneration and loss have been associated with ARHL [53,59]. In this regard, the reported increase in lumen size may be indicative of extensive capillary wall damage, including merging of luminal and perivascular spaces [55,59].

Thus, structural changes with aging in the microcirculation of the SV seem to be aggravated by central Aβ pathology and may contribute to worsen “naturally occurring” ARHL.

## 5. Conclusions

Compared with early, naturally occurring ARHL in the C57BL/6J mouse, in the *App*^NL-F^ mouse key ARHL traits further accelerate and increase in severity. These include larger threshold elevations starting at the adult age, lower rate of outer hair cell and spiral ganglion neuron survival, larger atrophy of the SV and damage to accompanying capillary blood vessels. Because no other phenotypes besides Aβ pathology seem to exist in the *App*^NL-F^ mice, the reported data support that central Aβ pathology linked to Alzheimer disease aggravates ARHL. As amyloid plaque accumulation is not evident in the cochlea during Alzheimer disease, other pathophysiological mechanisms must be explored. Increased damage to the microcirculation in the stria vascularis, propagated from alterations in central microcirculation is one possibility. Another possibility is potentiation by Aβ pathology of aging damage to descending corticofugal pathways modulating cochlear activity, known to participate in cochlear aging. These and other possibilities will require testing to consolidate the emerging notion of a vicious circle between ARHL and AD, with implications for its handling in the human clinic.

## Declarations

### Ethical Statement

#### Ethical approval statement for animal experiments

The use, care and handling of the animals followed current EU (Directive 2010/63/EU) and Spaińs (Law 32/2007; R.D. 53/2013) regulations. Procedures and protocols were approved by the competent authority (Dirección General de Ordenación Agropecuaria, Consejería de Agricultura, Ganadería y Desarrollo Rural, Gobierno de Castilla-La Mancha) with authorization Code ES020030000490 and supervised and monitored by UCLM’s Animal Experimentation Service in compliance with ARRIVE guidelines.

#### Data availability statement

The raw data supporting the conclusions of this article will be made available by the authors, without undue reservation.

#### Author contributions (CRediT)

Conceptualization: JMJ, JCA, VF-S. Data curation: JCA, VF-S. Formal analysis: JCA, VF-S, Z B, JMJ. Funding acquisition: JMJ, V F-S, JCA. TL. Investigation: VF-S, JCA, ZB, MCG-U, JMJ, TCS, TS. Methodology: VF-S, JCA, MCG-U, ZB. Project administration: JMJ. Resources: JMJ, TCS, TS. Software: JCA. Supervision: JMJ. Validation: All authors. Writing-original draft: JMJ, JCA, VF-S. Writing-review and editing: All authors.

## Acknowledgments

The authors express their appreciation to Prof. Dr. Carlos Dotti (CBM “Severo Ochoa”-UNAM) for his unvaluable assistance and advice in establishing the *App*^NL-F^ mice colony at UCLM.

## Abbreviations

Aβ: amyloid
AD: Alzheimeŕs disease
ANOVA: analysis of variance
APP: amyloid precursor protein
ARHL: age-related hearing loss
C57: C57BL/6J mouse
DAPI: 4′,6-diamidino-2-fenilindol
H/E: hematoxylin-eosin
KI: gene knock-in
OHC: outer hair cells
PF: paraformaldehyde
SD: standard deviation
SEM: standard error of the mean
SG: spiral ganglion
SPL: sound pressure level
SV: stria vascularis
WT: wild type

